# Cortico-striatal activity driving compulsive reward seeking

**DOI:** 10.1101/789495

**Authors:** Masaya Harada, Vincent Pascoli, Agnes Hiver, Jérôme Flakowski, Christian Lüscher

**Affiliations:** Department of Basic Neurosciences, Faculty of Medicine, University of Geneva, Geneva, Switzerland; Clinic of Neurology, Department of Clinical Neurosciences, Geneva University Hospital, Geneva, Switzerland

## Abstract

Addicted individuals compulsively seek drugs. Cortico-striatal projections have been implicated in persevering to seek rewards even when punished. The temporo-spatial determinants of the activity underlying the compulsive reward seeking however remains elusive. Here we trained mice in a seek-take chain, rewarded by optogenetic dopamine neuron self-stimulation (oDASS). Mice that persevered when seeking was punished, exhibited an increased AMPA/NMDA ratio selectively at orbitofrontal cortex (OFC) to dorsal striatum (DS) synapses. In addition, an activity peak of spiny projection neurons (SPNs) in the DS at the moment of signalled reward availability was detected. Chemogenetic inhibition of OFC neurons curbed the activity peak and reduced punished reward seeking, as did optogenetic hyperpolarization of SPNs time locked to the cue predicting reward availability, establishing a causal link. Taken together, we conclude that the strengthening of OFC-DS synapses drives SPNs activity when a reward predictive cue is delivered, thus encouraging reward seeking in subsequent trials.

## Introduction

Compulsive drug seeking is a hallmark of addiction ^1^. In humans this behaviour has deleterious consequences on physical, social and economic well-being. While addictive drugs are widely used recreationally, only a small fraction of individuals lose control over drug intake ^2^. The individual difference of the vulnerability to addiction has also been observed in laboratory animals, where some individuals continue to self-administer drug rewards even when facing a punishment while others cease drug taking ^3–5^. These models apply punishment as a noxious electrical foot shock ^6^ or in the form of aversive, bitter taste for oral self-administration ^7^. Here we define compulsion as the perseverance of self-administration in light of punishment. Across different drug self-administration schedules and with multiple classes of addictive drugs, including psychostimulants and alcohol compulsion is observed in a fraction of animals ^3,7–9^. Long lasting exposure and escalation of drug intake are prerequisites for compulsive use ^3^.

Here we focus on compulsive drug seeking, which may involve distinct neural substrates and cognitive strategies compared to compulsive drug-taking behaviour ^6^. Compulsive drug seeking may reflect the incapacity to shift from habitual to goal-directed motivational control ^10^ while compulsive drug taking could be explained by excessively narrow goal directed behavior ^1^. In rodents compulsive drug seeking can be modeled by using a seek-take chain schedule of cocaine self-administration that includes a random time interval, after which the animal has to press a seeking lever to gain access to the taking lever. Once fully acquired, an aversive stimulus is delivered to the seeking lever instead of the taking lever in a fraction of trials.

During early stage of addiction, drug-evoked dopamine (DA) transients in the nucleus accumbens (NAc) positively reinforce self-administration ^11,12^. Resulting dopamine-dependent synaptic plasticity in the NAc and the ventral tegmental area (VTA) have been causally linked to drug adaptive behaviours, including locomotor sensitization and cue associated seeking behaviour ^13–17^. At later stages of drug exposure, transition to compulsive drug use may be mediated by the recruitment of potentiated synapses in more dorsal areas of the striatum ^1,18^. The dorsal striatum (DS), which is also the site of habit formation ^19,20^, may therefore play a role in the incapacity to shift from habitual to goal-directed behaviour, which defines compulsive drug seeking. Consistent with this hypothesis, pharmacological inhibition of the dorsolateral striatum reduces compulsive drug seeking behaviour ^21^ as well as habitual drug seeking ^20^.

Top-down control of the striatum arises in the anterior cortex, where neurons of the medial prefrontal cortex (mPFC), orbitofrontal cortex (OFC) and motor cortex (M1) send their axon to subregions of the DS. The striatum can be separated into the dorsolateral, dorsomedial, and ventral striatal subdivisions, defined by their topographic organization and relative functional involvement in sensorimotor, associative, and limbic process, respectively ^22–27^. A recent study recapitulated the connectome of three main projections in the DS ^28^. mPFC to dorsomedial, OFC to central and M1 to dorsolateral striatum may all be implicated in compulsive reward seeking, but their respective role remains elusive. Contrasting mPFC hypofunction and OFC hyperfunction may both favour the appearance of compulsion ^5,29–31^. Motor cortex, believed to control habitual responding via its projection to the dorsolateral striatum, may be involved in compulsive drug seeking, if there is a failure to disengage habitual behaviour ^32^.

Although several cortico-striatal pathways have been implicated in compulsive drug seeking, it is not known whether pathway-specific activity patterns and synaptic plasticity emerging after drug self-administration are critical to reinforce drug seeking despite the risk of negative consequences. To tackle these questions, we used optogenetic activation of VTA DA neurons model to replace drugs in which mice self-stimulate in a seek-take chained schedule. Since all addictive drugs induce their reinforcing effects by elevating DA in the NAc ^11,12^, this optogenetic stimulation allows us to focus on mechanisms common to several addictive drugs while avoiding off target effects of individuals substances. Moreover, optogenetic self-stimulation of VTA DA neurons evokes analogous plasticity to cocaine in the VTA ^33^ and nucleus accumbens ^5^, and results in a dissociation between punishment-sensitive and punishment-resistant phenotypes. To probe for compulsive reward seeking, we tested whether mice would persevere in the seek-take chain despite having to endure a foot-shock. We found that this behavior was bimodally distributed with about 60% of the mice fulfilling the criterion for compulsive reward seeking. Combining circuit tracing, *ex vivo* assessment of synaptic transmission, photometry and chemogenetic as well as optogenetic manipulations, we revealed a critical involvement of OFC-DS projections in compulsive seeking behaviour. We parse the activity peak in SPN activity at a specific moment of the seek-take chain that leads the animal to engage in subsequent trials, thus providing a cellular explanation for compulsive seeking.

## Results

### Compulsive oDASS seeking in a fraction of mice

We injected the VTA of DAT-Cre mice with an AAV5-EF1a-DIO-channelrhodopsin-2 (ChR2)-eYFP and implanted an optic fiber just above the injection site (Fig. 1a). Two weeks later, mice first acquired a taking response and then were trained under a seek-take chain (Fig. 1b-d), where they had to press the seek lever to access the take lever (see methods for details). All mice completed 30 trials (random interval; RI = 60s) in 4 daily baseline sessions (Fig. 1d,e). The mice then underwent 5 days of punished sessions, during which a foot-shock (0.25 mA, 500 ms) was delivered with the first press on the seek lever after the end of the RI in 30% of the trials, while in 70 % of the trials the take lever was presented (Fig. 1b,c). This schedule caused some mice to stop pressing on the seeking lever (Fig. 1e and Extended Data Fig.1 and 2). Using unsupervised classification method on five behavioral parameters (session duration, number of seek lever press, completed trials, delay for the first seeking lever press, and delay for the last seeking lever press), two distinct groups of mice emerged (Fig. 1f and Extended Data Fig. 2). About 40 % strongly reduced their seeking behaviour and were unable to complete most of the trials under punishment, which we called “renouncers”. The second group, here called “perseverers”, about 60% of all mice, maintained sufficient number of seeking lever presses under punishment to complete most of the trials (Fig. 1e and Extended Data Fig. 2). The number of seek lever presses during the baseline sessions was not different between the two groups and could thus not predict perseverance under punishment (Fig. 1g and Extended Data Fig. 2). Overall, the fraction of completed trials was significantly reduced during punished sessions in renouncers but not in perseverers. We defined the latter as exhibiting compulsive seeking behavior.

**Fig. 1.**
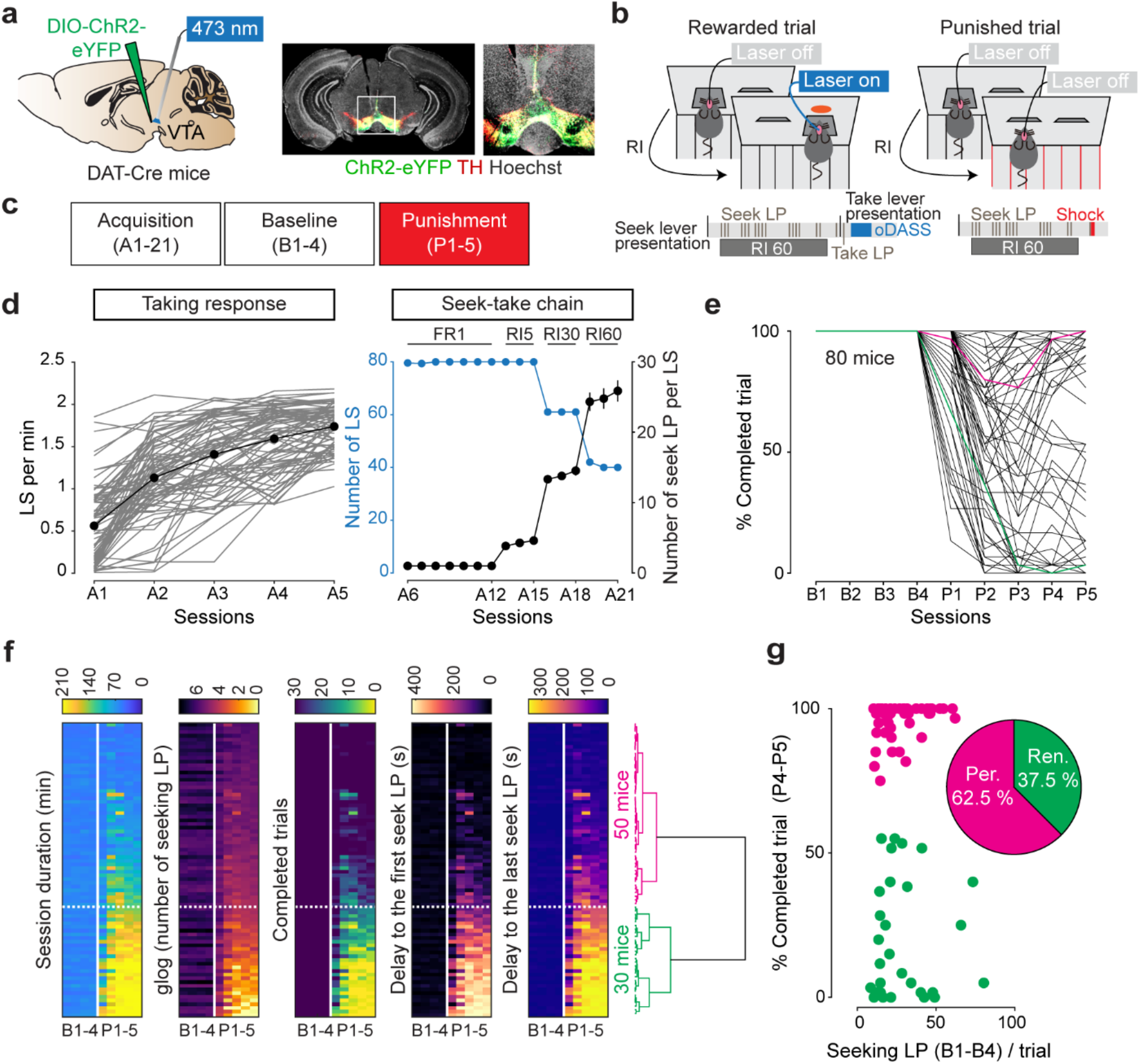
oDASS seeking despite negative consequences in a subset of mice. **a,** Schematic of optic fibre implantation into the VTA of DAT-Cre mice (left), infected with AAV5-EF1α-DIO-ChR2-eYEP and staining of tyrosine hydroxylase (middle and right). **b,** During rewarded trials, the first seek lever press after the end of the RI triggered the retraction of the seek lever and the presentation of the take lever. Single press on the take lever resulted in the delivery of the laser stimulation. During punished trials, the first seek lever press after the end of RI triggered the retraction of the seek lever and the delivery the foot shock (500 ms, 0.25 mA, right). **c**, Experimental timeline. **d,** Acquisition of seek-take chained schedule for oDASS. (A1-21). During taking response training, only the taking lever was presented with the seeking lever retracted. Single press on the taking lever triggered the laser stimulation (left). After 5 sessions of taking response training, acquisition sessions of seek-take chained schedule started (right). **e,** Percent of completed trials during baseline (4 d) and punished (5d) sessions. Green and pink lines show the examples of two mice in Extended data Fig. 1. During baseline sessions, all the trials were rewarded. During punishment sessions, 70% of trials were still rewarded and the other 30% of the trials were punished. **f,** Unsupervised clustering method yields two clusters (Renoucer and Perseverer). **g,** Percent of completed trials as a function of number of seek lever presses *per* trial during baseline sessions.

### Parallel cortico-striatum streams

To probe the underlying neural circuits, we characterized cortico-striatal pathways, which connect specific cortical areas to sub-regions of the DS (Hunnicutt et al., 2016). We injected the retrograde tracers CTB of distinct colours into medial, central and lateral parts of the dorsal striatum of the same mouse (mDS,cDS and lDS, respectively. Fig. 2a,b). With the CTB-488 seeded in the mDS, we observed neurons retrogradely labelled predominantly in the mPFC, especially within the prelimbic cortex (Fig. 2c,d). By contrast, the CTB-555 injected in the central part of the dorsal striatum retrogradely labelled neurons in the OFC (Fig. 2c,d). With the CTB-647 injected in the lateral DS, cells in the primary motor cortex (M1) were stained (Fig. 2c,d). These data indicate that sub-regions of the DS receive inputs from distinct cortical regions with only very sparse mixed innervation (less than 20%). To probe synaptic transmission, we next injected the anterograde tracer AAV8-hSyn-ChrimsonR-tdTomato in mPFC, OFC or M1 (Fig. 3a) yielding fibers in the respective subregions of the DS (Fig. 3b). Selective optogenetic activation in acute brain slices produced large light-evoked EPSCs in SPNs of the mDS, cDS and lDS respectively (Fig. 3c), again with only minimal mixed inputs found in DS subregions. Taken together our functional and anatomical investigations confirm the existence of largely non-overlapping cortico-striatal information streams.

**Fig. 2.**
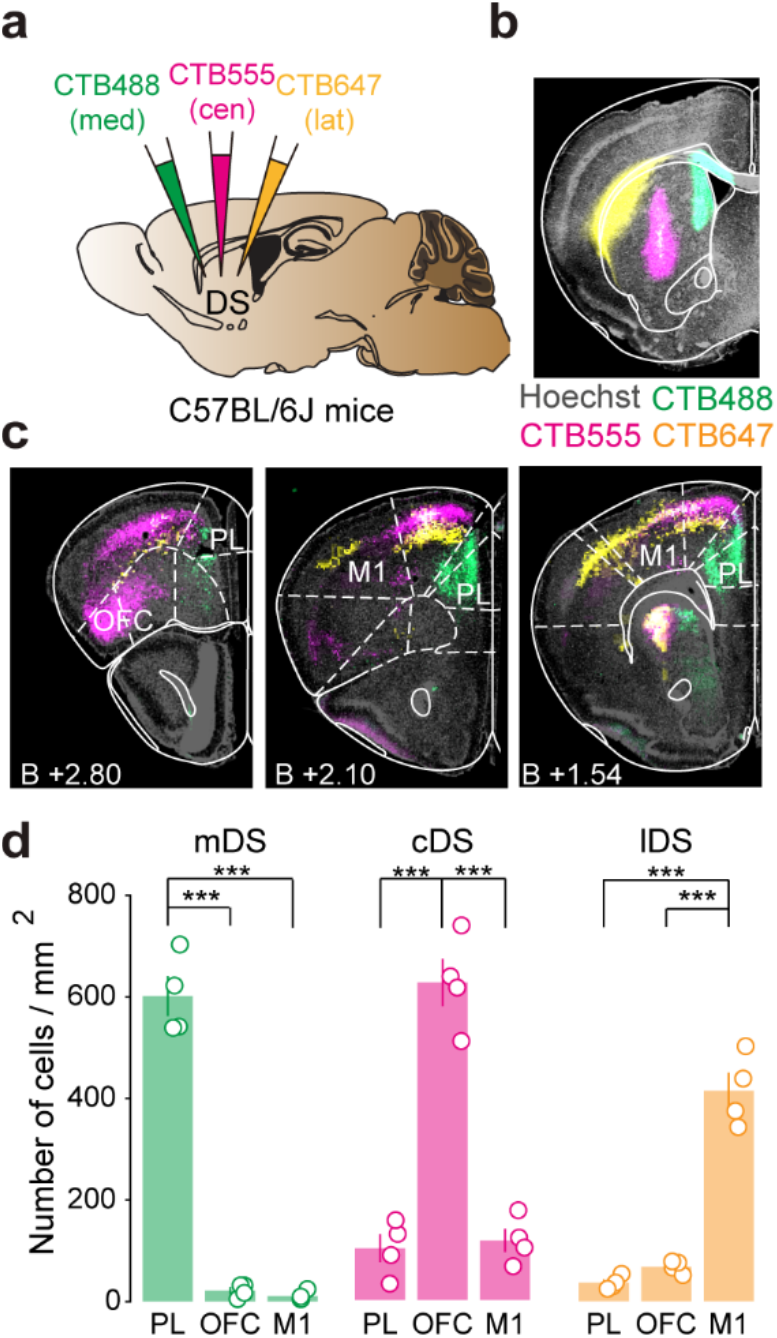
Three parallel cortico-striatal pathways. **a,** Schematic for the experiment. **b,** Retrograde tracer, CTB conjugated to the dye Alexa 488 (green), 555(pink), or 647 (yellow) were injected in medial, central or lateral part of the dorsal striatum (DS), respectively. **c,** Example pictures taken from cortex. **d,** Quantification in Prelimbic cortex (PL), orbitofrontal cortex (OFC) and primary motor cortex (M1) for each striatal projection. Projections to medial part of the DS (mDS, left). F(2,9)=215.9, p<0.0001, Post-hoc comparison with Bonferroni test yielded: p<0.0001 for PL versus OFC; p<0.0001 for PL versus M1. Projections to central part of the DS (middle). F(2,9)=76.98, p<0.0001, Post-hoc comparison with Bonferroni test yielded: p<0.0001 for PL versus OHC; p<0.0001 for OFC versus M1. Projections to lateral part of the DS (right). F(2,9)=99.75, p<0.0001, Post-hoc comparison with Bonferroni test yielded: p<0.0001 for PL versus M1; p<0.0001 for OFC versus M1 Error bars, s.e.m. ***p < 0.001

**Fig. 3.**
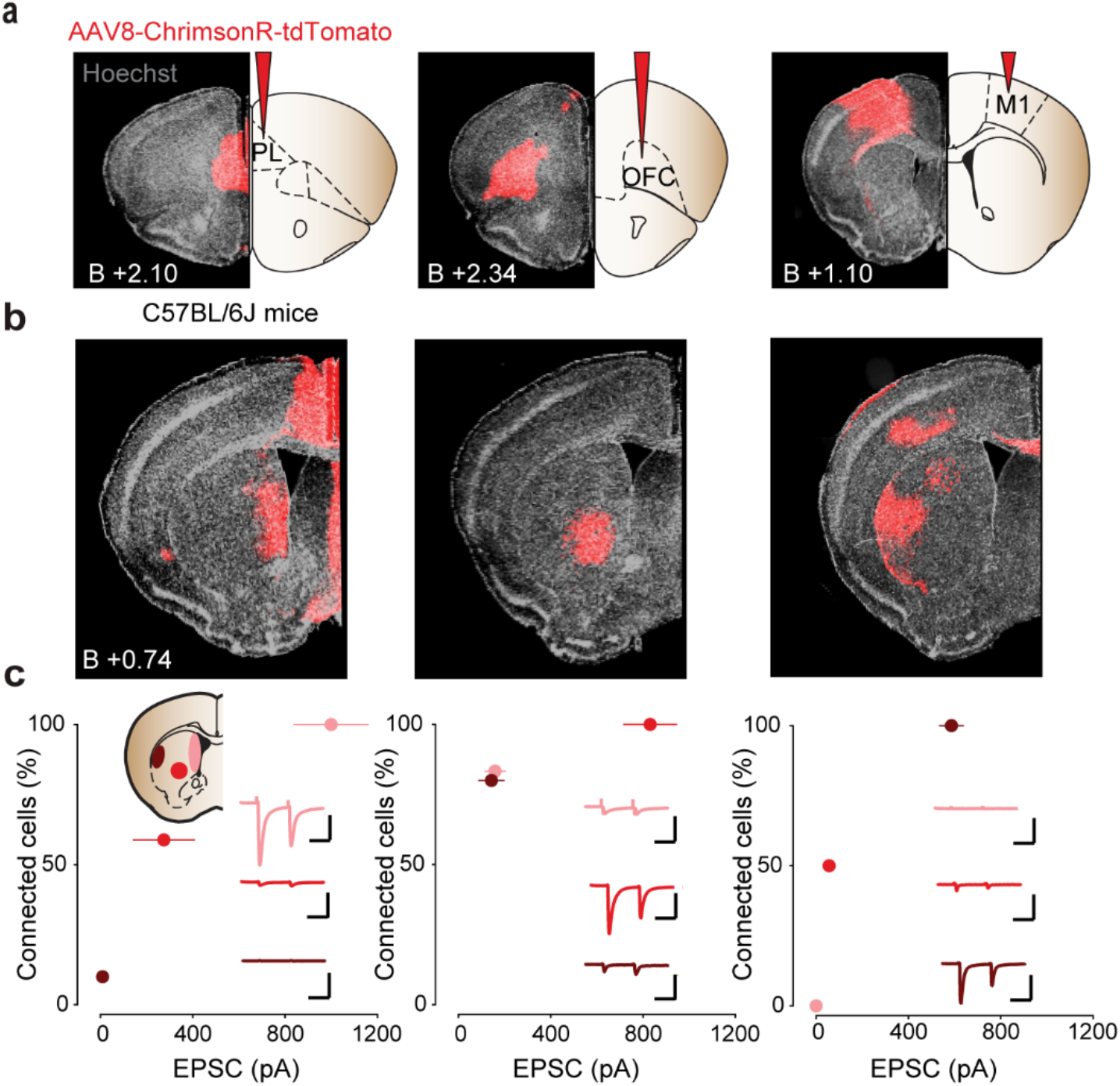
Functional connectivity at three parallel cortico-striatal pathways. **a,** Coronal images of mice brain slices infected with AAV8-hSyn-ChrimsonR-tdTomato in the prelimbic cortex (PL, left), orbitofrontal cortex (OFC, centre) and primary motor cortex (M1, right). **b,** Coronal images of the dorsal striatum corresponding to the images of panel **a**. **c,** Functional connectivity between PL (left) OFC (middle), and M1 (left) and DS subregions. EPSCs were recorded in medial (light red), central (red) and lateral (dark red) part of the striatum. Number of recorded neurons: medial/central/lateral, PL: 12/17/10, OFC: 12/12/10, M1: 12/12/12. Scale bars. 50ms, 500pA. Error bars, s.e.m.

### Selective plasticity of cortico-striatal synapses in compulsive mice

Next, we assessed the synaptic strength of the three cortico-striatal streams *ex vivo* in mice that underwent punished oDASS seeking. To minimize interference with oDASS, we expressed the red-shifted excitatory opsin ChrimsonR either in the mPFC, OFC or M1 of DAT-cre mice infected with ChR2 in the VTA (Fig. 4a,c,e) and prepared acute slices 24h after the last punished session. Since previous results did not show any difference between D1R and D2R SPNs in synaptic strength in the dorsal striatum ^29^, and both populations are activated by acute cocaine injection ^34^, these experiments were done without SPN distinction. To quantify the synaptic strength at the three synapses, we measured the AMPAR-EPSCs to NMDAR-EPSCs ratio (A/N) at positive potential (+40mV, with AP5 pharmacological isolation of AMPAR-EPSCs and calculation of NMDAR-EPSCs) and used naïve mice as controls. We found that the A/N at OFC-cDS was higher in slices from persevering mice than in slices from renouncing or naïve mice (Fig. 4e) while at mPFC-mDS and M1-lDS synapses, the A/N in persevering, renouncing or naïve mice were not different (Fig. 4c,f). Of note, the A/N ratio was smaller at M1-lDS than at the other synapses in all three groups of mice. Taken together, these data demonstrate that synaptic strength was selectively increased in compulsive mice at OFC-cDS. We also measured rectification index at these three cortico-striatal pathways, detecting no differences between groups (Extended Data Fig. 3).

**Fig. 4.**
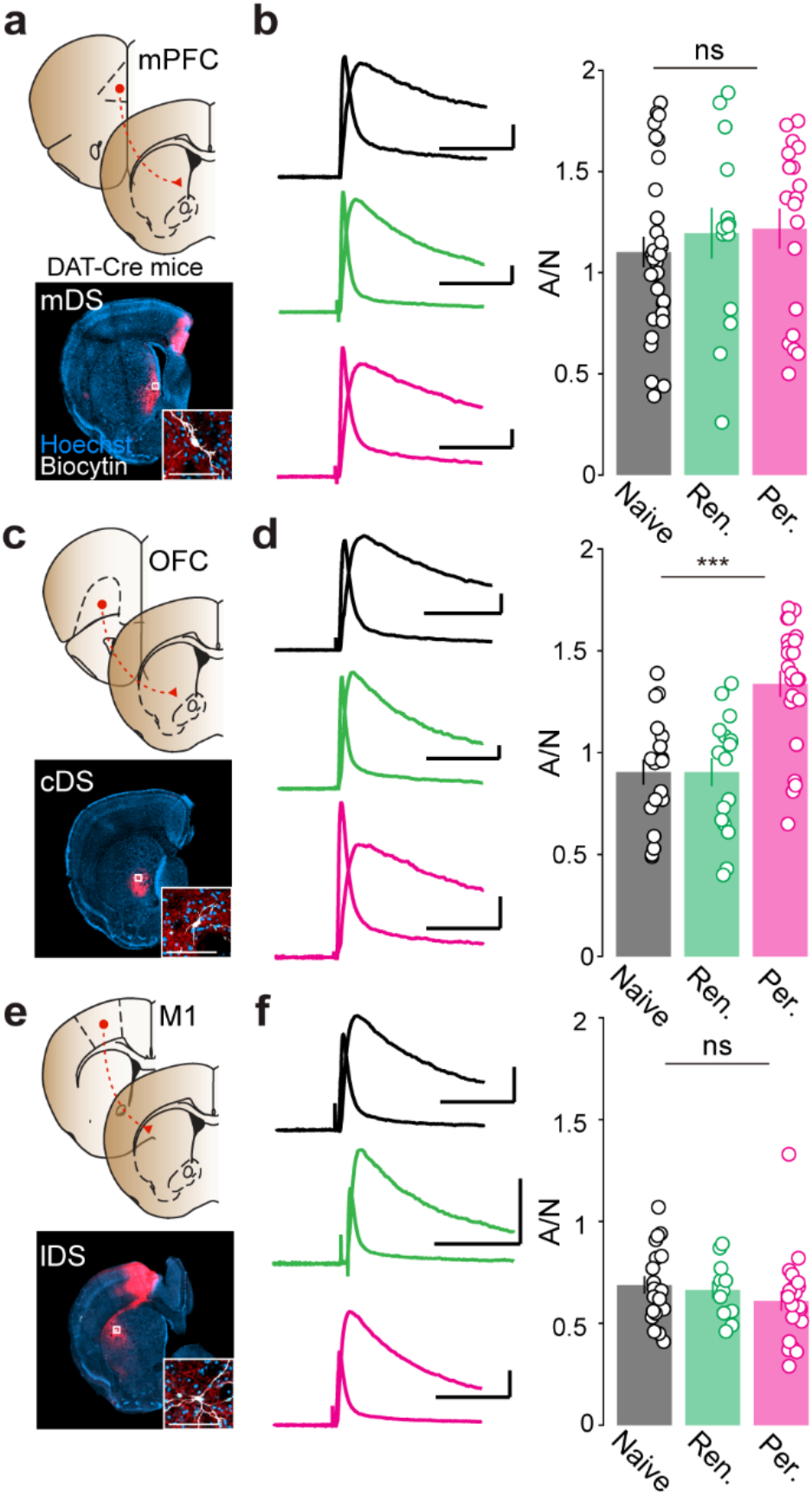
Specific potentiation at OFC-cDS synapses in persevering mice. **a,** Schematic of the preparation for ex vivo recordings at medial prefrontal cortex (mPFC)–medial part of the dorsal striatum (mDS) synapses (top). Coronal section shows terminals in the striatum and recorded neuron (bottom). Inset shows recorded neuron at high magnification. **b,** Example traces (average of 20 sweeps) of AMPA and NMDA-EPSCs, recorded at +40 mV in mDS slices from naive, renouncer or perseverer mice (left). A/N of three groups are not statistically different (right) (n=31/14/19, naive/renouncer/perseverer, F(2,61)=0.4941, P=0.62). **c,** Schematic of the preparation for **d**(top). Coronal section shows terminals in the striatum and recorded neuron (bottom). Inset shows recorded neuron at high magnification. **d,** Example traces (average of 20 sweeps) of AMPA and NMDA-EPSCs, recorded at +40 mV in cDS slices from naive, renouncer or perseverer mice (left). A/N is increased in perseverer mice but not renouncer mice (right) (n=19/17/23, naive/renouncer/perseverer. F(2,56)=15.95, p<0.0001, Post-hoc comparison with Bonferroni test yielded: p<0.0001 for naive versus perseverer; p<0.0001 for renouncer versus perseverer. **e,** Schematic of the preparation for **f**(top). Coronal section shows terminals in the striatum and recorded neuron (bottom). Inset shows recorded neuron at high magnification. **f,** Example traces (average of 20 sweeps) of AMPA and NMDA-EPSCs, recorded at +40 mV in lDS slices from naive, renouncer or perseverer mice (left). A/N of three groups are not statistically different (right) (n=17/11/21, naive/renouncer, F(2,46)=1.23, p=0.30).Scale bars, 50 ms, 200 pA, and 100 μm. Error bars, s.e.m. ***p < 0.001

### Activity peak at seek lever retraction in persevering mice

We next asked when the synaptic potentiation altered neural activity in compulsive mice *in vivo*. To this end, we monitored intracellular calcium as a *proxy* for neural activity, expressing AAV5-CamKII-GCamp6f in the cDS and recording fluorescence with an optic fiber (Fig. 5a). To spectrally separate the calcium signal from oDASS, we used AAV8-hSyn-DIO-ChrimsonR-tdTomato for the self-stimulation (Fig. 5a). During baseline sessions, a robust calcium peak was detected after the seek lever retraction in both perseverers and renouncers (Fig. 5b,c), albeit with an amplitude that was higher in mice later identified as perseverers (Fig. 5c). Aligned to seek lever extension, seek lever press or take lever press, only much smaller increase in calcium signal could be detected (Extended Data Fig. 4). Take lever was presented 0.5 s after the seek lever retraction (Fig. 5b); to determine whether this signal is mostly due to the seek lever retraction or contaminated by the presentation of the taking lever, in a separate experiment, the taking lever was extended with a 3s delay (Extended Data Fig. 5). Using this delayed interval to associate the behavioral events confirmed that peak calcium activity coincided primarily with the retraction of the seeking lever, with only a small contribution of the subsequent take lever presentation during unpunished sessions (Extended Data Fig. 5). During punishment sessions, the calcium signal at seek lever retraction was still observed in persevering mice but disappeared in renouncers (Fig. 5d). To compare this reduction in the 2 experimental groups, the peak amplitude around seek lever retraction during punishment session was normalized to the peak amplitude during baseline sessions (Fig. 5e). This confirmed the positive correlation between the conservation of SPN activity and compulsion (Fig. 5f). By contrast, in the lateral part of the dorsal striatum no significant difference of the calcium signals between perseverers and renouncers around seek lever retraction (Extended Data Fig. 6), seek lever presentation, seek lever press or take lever press (Extended Data Fig. 7) was observed. We did not observe any difference in normalized amplitude around seek lever retraction (Extended Data Fig. 6), indicating that the amplitude did not change depending on the number of the trials. In sum, the retraction of the seek lever caused a peak of activity, which was sustained during punishment sessions in perseverers, while in renouncers the calcium signal declined in amplitude during punishment sessions. This observation was specific to the central part of the DS, the preferred target regions of neurons located in the OFC.

**Fig. 5.**
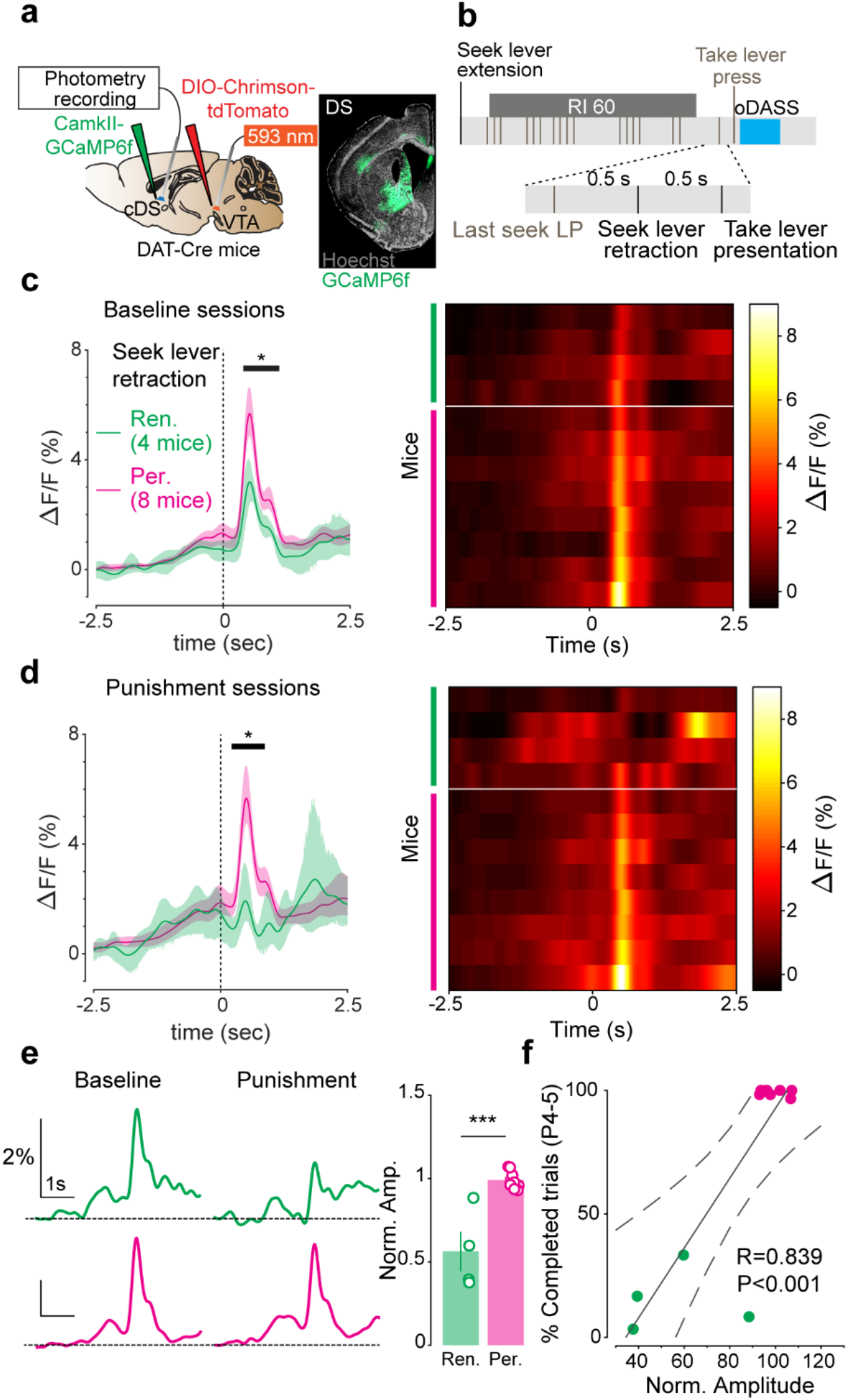
Persistent hyperactivity in cDS around seek lever retraction in perseverer mice. **a,** Schematic of the preparation for in vivo photometry recording (left), and coronal image of a mouse brain slice infected with AAV5-CamkII-GCaMP6f in cDS (right). **b,** Schematic for the behavioural paradigm. **c,** Calcium signal (ΔF/F) around the seek lever retraction during baseline sessions for persevering and renouncing mice (left). Black line and asterisk indicate time window with significant difference between perseverer and renouncer and shaded area represents 95% confidence interval. Significant difference at 0.4361 to 1.0949 s. The heatmap shows calcium signal for individual mouse (right). **d,** Calcium signal (ΔF/F) around the seek lever retraction during punishment sessions for persevering and renouncing mice (left). Black line and asterisk indicate time window with significant difference between perseverer and renouncer and shaded area represents 95% confidence interval. Significant difference at 0.2198 to 0.8687 s. The heatmap shows calcium signal for individual mouse (right). **e,** Examples of calcium signals for renouncer (green) and perseverer (pink) around the seek lever retraction during baseline and punishment sessions (left). Peak amplitude during punishment sessions is normalized to the amplitude during baseline sessions. Renouncers showed significant reduction of peak amplitude (right) (P= 0.0005,t_10_ =5.071). **f,** Percent of completed trials (P4-5) as a function of normalized amplitude of calcium signals around the seek lever retraction. Percent of completed trials is positively correlated to normalized amplitude (Pearson correlation coefficient (R)=0.839, P<0.001). Error bars, s.e.m. *p < 0.05; ***p < 0.001.

### OFC inhibition during compulsive oDASS seeking

To test for the contribution of OFC-cDS projections in the activity peak of SPNs of the cDS at seek lever retraction, we expressed the inhibitory DREADD (designer receptors exclusively activated by designer drugs: CamKIIa-hM4D) in excitatory neurons of the OFC (Fig. 6a,c). To validate the chemogenetic approach, we evoked OFC-DS transmission and bath-applied CNO (clozapine-N-oxide, 10μM) to acute DS slices of mice infected with both hM4D and chrimson in the OFC. CNO reduced optogentically evoked-EPSCs recorded from SPNs (Fig. 6b), most likely by preventing glutamate release from OFC terminals ^35^. Next, to silence the OFC during behavior, we injected AAV1-CamkII-hM4D-mCherry in the OFC and AAV8-hSyn-DIO-ChrimsonR-tdTomato in the VTA of DAT-cre mice. In addition, AAV5-CamKII-GCamp6f was injected in the cDS and an optic fiber was positioned to allow photometry recording of SPNs activity with or without OFC chemogenetic inhibition (Fig. 6c). CNO injection (2mg/kg, IP) 1h before the recording session significantly attenuated the calcium peak at seek lever retraction (Fig. 6d). This experiment was conducted only in persevering mice previously selected during additional punished sessions. Finally, chemogenetic inhibition of OFC of persevering mice was sufficient to reduce the number of completed trials during punished sessions (Fig. 6e). The relative reduction of completed trials was proportionate to the attenuation of the calcium peak (Fig. 6f). Histological verification indicated the expression of CamkII-hM4D-mCherry in all mice (Extended Data Fig. 8). Taken together these data indicate that activity of SPNs at seeking lever retraction was driven by excitatory inputs form the OFC, which, when inhibited, reduced compulsive seeking for oDASS.

**Fig. 6.**
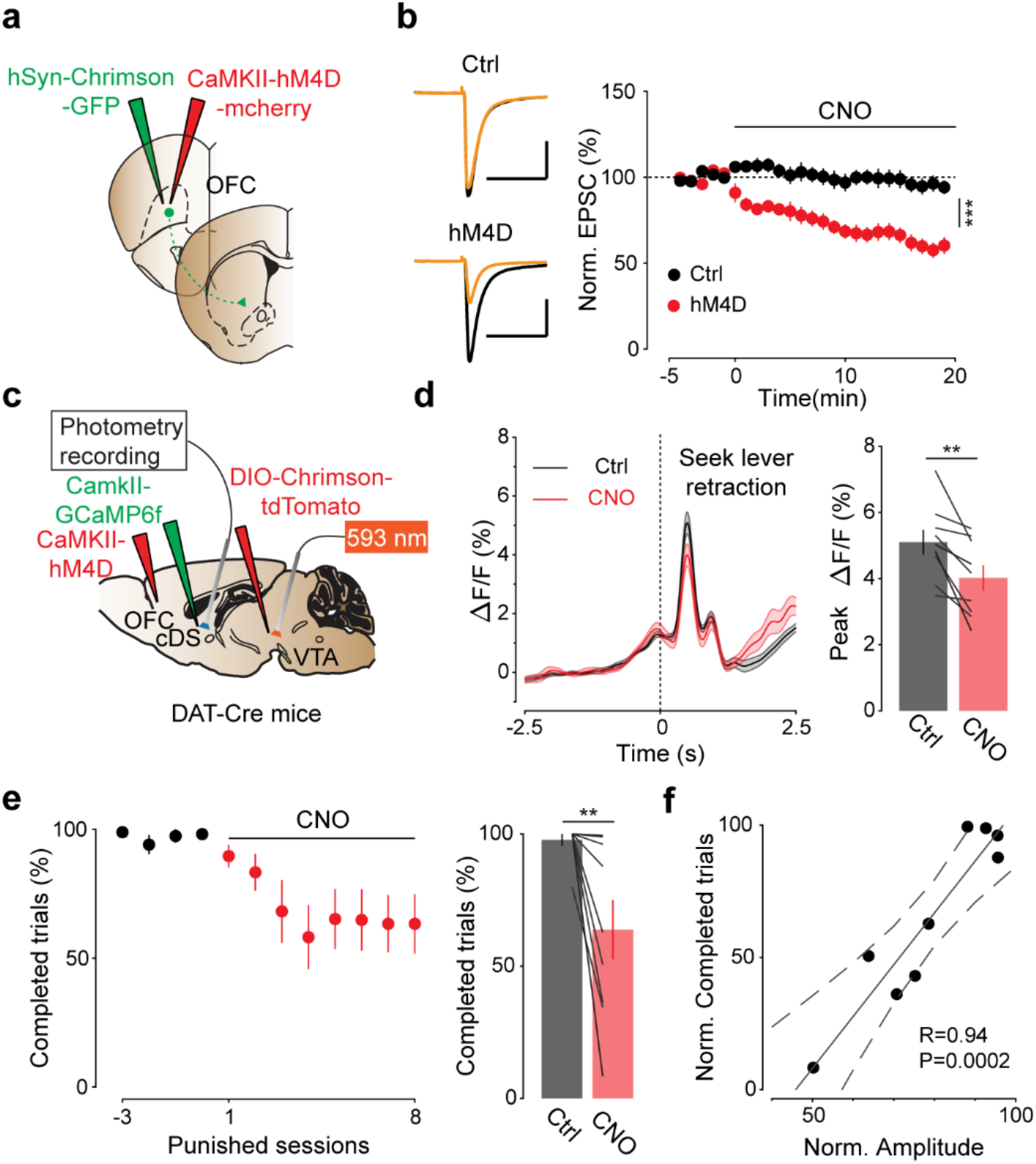
Attenuation of compulsive reward seeking and flattening of calcium signal around seek lever retraction by chemogenetic inhibition of the OFC. **a,** Schematic for the preparation. **b,** Example traces of AMPA EPSCs recorded at −70 mV from control mice (top, left) and mice expressing hM4D in the orbitofrontal cortex (OFC, bottom, left) during baseline (black) and 15 min after clozapine N-oxide (CNO, 10 μM) bath application (yellow). Control mice received injection of AAV9-hSyn-ChrimsonR-GFP but not AAV1-CamKII-hM4D-mcherry. Group data for normalized EPSCs as a function of time (middle). CNO decreased light evoked EPSCs in the central part of the dorsal striatum (cDS).t_20_ =5.049, P< 0.0001, Scale bars, 50ms, 500pA. Number of recorded neurons, Ctrl/hM4D; 10/12. **c,** Schematic for the preparation. **d,** Calcium signals around seek lever retraction before CNO injection (black) and during CNO injection (red, left). Shaded area represents s.e.m. CNO injection reduced peak amplitude (t_8_=3.926, P= 0.0044, right). **e,** Percent of completed trials over sessions (left). After 4 sessions of punishment, CNO was injected 1 hour before each session (2mg/kg, IP). Percent of completed trials on the last 2 days before CNO injection and the last 6 days during CNO injections (right). CNO injection reduced percent of completed trials (t_8_=3.185, P= 0.0129). **f,** Normalized calcium signal amplitude (before CNO/ during CNO) as a function of normalized completed trials (before CNO/ during CNO). Two parameters are positively correlated (Pearson correlation coefficient (R)=0.938, P=0.00019). Error bars, s.e.m. **p < 0.01; ***p < 0.001

### Time locked inhibition of SPN activity

To probe the exact timing of the OFC-DS activity required, we next inhibited SPNs time locked to the seek lever retraction. To this end, we injected AAV5-CamKII-eArchT3.0-eYFP in the cDS and placed optic fibers bilaterally (Fig. 7a,b). In acute slices, orange light (wavelength; 585 nm) activated ArchT3.0 and effectively suppressed action potentials induced by a depolarizing current injection (Fig. 7c). First, mice underwent 5 sessions of punished oDASS to identify perseverers. Next, additional 9 punished sessions were performed with or without inhibition of cDS SPNs (shown as test and ctrl, respectively) for 4s starting at the first seek lever press after the end of RI to silence SPNs during the seek lever retraction (Fig. 7d, Top). Compulsive seeking parameters were reduced by the time-locked cDS inhibition but recovered quickly in sessions without inhibition (Fig. 7d,f). Importantly, this inhibition at seek lever press did not affect seeking in baseline conditions (Extended Data Fig. 8). Moreover, applying optogenetic inhibition at the first seek lever press of each trial (Fig. 7e, top), when only small calcium transients were observed (Extended Data Fig. 4), had no impact on compulsive seeking behavior (Fig. 7e,f). We conclude that brief cDS inhibition is not aversive per se, but can specifically reduce seeking behavior selectively in perseverer mice.

**Fig. 7.**
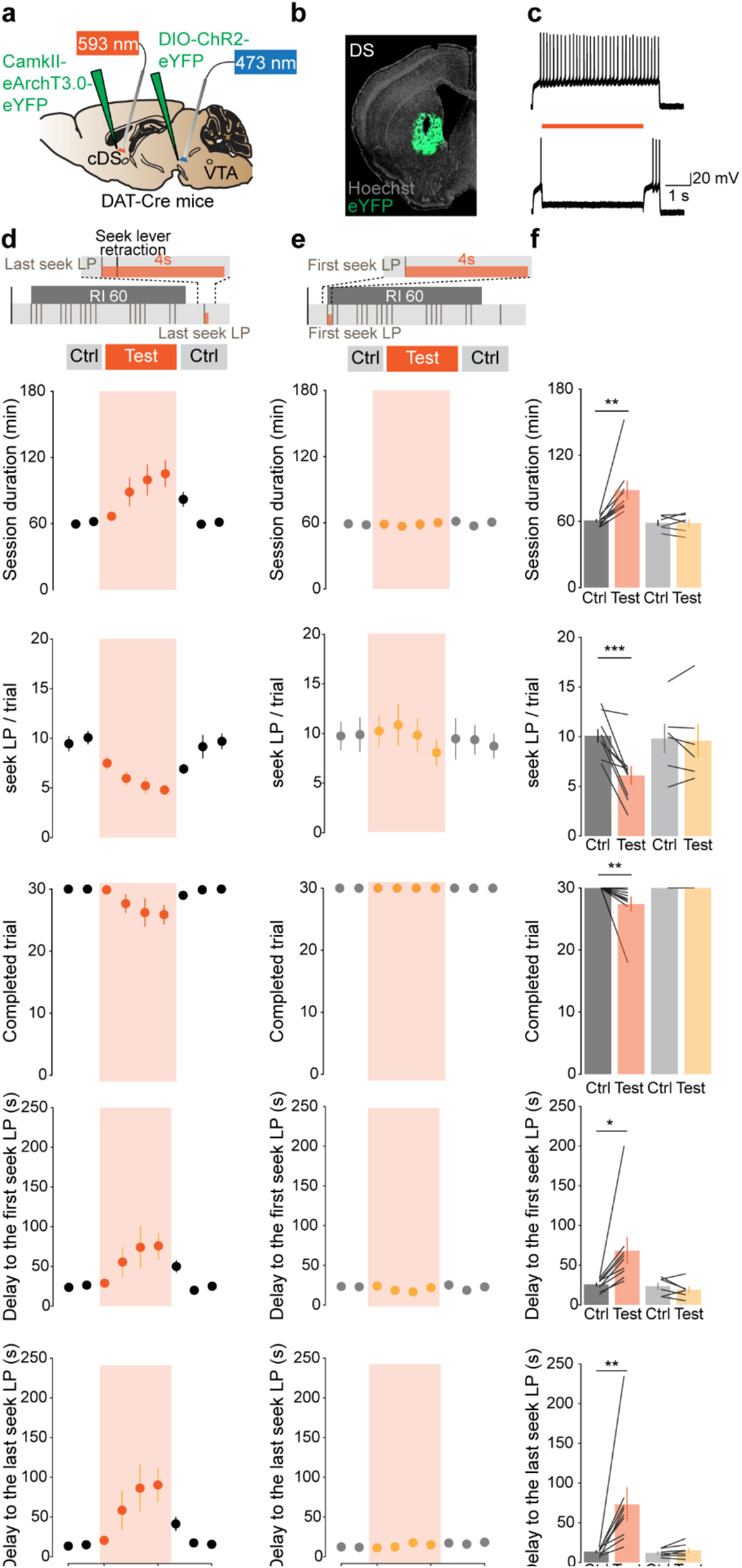
Attenuation of compulsive reward seeking by time locked inhibition in the cDS. **a,** Schematic for the preparation. **b,** Image of a mouse brain infected with eArchT3.0-eYFP in the cDS. **c,** Orange light (593 nm) suppressed action potentials induced by current injections (300pA, 5s) in eArchT3.0 expressing neurons. **d,** Schematic for the inhibition protocol (Top). Five parameters used for the clustering in Fig. 1 **f**(bottom). **e,** Schematic for the inhibition protocol (Top). The same parameters as in **d**(bottom). **f,** Group data for panel **d** and **e**. Behaviour was modified by the inhibition at the moment of the last seek LP but not first seek LP. Average of two sessions before inhibition (session −1 and 0) and last three days of inhibition (session 2 to 4) are shown as Ctrl and Test, respectively. Session duration; mixed two-way ANOVA: laser timing, F(1,13)=6.542, P=0.0238; treatment, F(1, 13)=7.603,P=0.0163; interaction, F(1,13)=7.638, P=0.0161. Bonferroni post hoc analysis, **p<0.01. Number of seeking lever presses per trial; mixed two-way ANOVA: laser timing, F(1,13)=1.098, P=0.3138; treatment, F(1,13)=15.08, P=0.0019; interaction, F(1,13)=12.00, P=0.0042. Bonferroni post hoc analysis, ***p<0.001. Number of completed trials; Wilcoxon matched-pairs signed rank test. P value was adjusted with Bonferroni-Dunn method. *P<0.01. Delay to the first seeking lever press; mixed two-way ANOVA: laser timing, F(1,13)=4.538, P=0.0528; treatment, F(1,13)=3.696, P=0.0767; interaction, F(1,13)=5.469, P= 0.0360. Bonferroni post hoc analysis, ***p<0.05. Delay to the last seeking lever press; mixed two-way ANOVA: laser timing, F(1,13)=4.472, P=0.0543; treatment, F(1,13)=6.024, P=0.0290; interaction, F(1,13)=4.733,P= 0.0486. Bonferroni post hoc analysis, **p<0.01. Error bars, s.e.m.

## Discussion

Here we observed compulsive reward seeking in about 60% of the mice that underwent oDASS, a mouse model of addiction. We identified a selective synaptic potentiation at OFC to SPNs synapses in the DS, along with a peak in calcium activity in these DS SPNs at the moment of the seeking lever retraction as neural correlates of compulsive oDASS seeking. Chemogenetic or optogenetic inhibition of OFC and DS respectively established that the DS SPNs activity peak depends on excitatory input from the OFC and is required for compulsive seeking in subsequent trials.

Compulsive drug seeking in addiction is an extreme form of decision bias, where the subject continues to seek the drug reward despite the negative consequences ^1,10,36^. The OFC has been implicated in this behaviour, based on experimental observations of compulsive reward taking ^5,29^. This also seems plausible as one of the physiological role of the OFC is to calculate cost-benefit, or economic value ^37^.

Contrasting the hypothesis of a general loss of cortical control ^30,38^, we found that enhanced OFC to DS transmission is associated with compulsion ^29^. Such gain of function thus represents an appealing alternate to prevailing explanations. The enhanced activity in the DS, driven by the OFC after seeking lever press followed by its retraction (a signal of trial completion that advertises reward availability) may alter the perceived cost-benefit balance of the behaviour. The fact that the signal decreases under punishment risk in renouncers suggests a degradation of the cue value. It is therefore likely that the activity peak codes for the reward value rather than punishment severity. In this scenario, perseverers continue to ascribe increasing reward values to drug-predictive cues, while the perception of the punishment remains intact. Indeed, it has been reported that OFC encodes this integrated cue value ^39,40^ along with the prediction of cue outcomes ^41,42^. Moreover, we have previously shown that perseverers and renouncers react to an unrelated painful stimulus equally ^5^. Our present findings with reward seeking reveal a circuit that is largely overlapping with the circuit implicated in compulsive reward taking ^29^, underscoring the involvement of the OFC to DS pathway across core features of addiction. Similar to what has been observed with compulsive oDASS or drug taking ^29,43^, the bimodal distribution for punishment resistant oDASS-seeking develops even in the genetically identical background of inbred animals. Moreover, the fraction of animals showing compulsive seeking for optogenetic stimulation of VTA DA neurons (approximately 60%) using the seek-take chain schedule is very similar to what was pxsreviously determined with more simple version of oDASS compulsive taking task ^29^. All together, these data suggest that vulnerability to compulsive oDASS seeking and taking involve the overlapping neural substrate of potentiated OFC-to-cDS transmission.

We did not find alterations in mPFC to mDS, nor M1 to lDS projections. This does not preclude a concomitant role of the prelimbic in compulsion ^30^ because of reduced behavioural inhibition. Recently, it has been reported that prelimbic to NAc pathway is involved in compulsive reward taking ^31^. Afferents onto SPNs of the lDS, which is the major target of the projection from M1, also remained unchanged with our paradigm. Since the latter has been implicated in habitual responding in drug self-administration protocols, our findings do not provide direct evidence that compulsive seeking arises from failure to disengage from habitual performance ^32^.

In the present study, we used compulsive oDASS seeking as an addiction model. This has the advantage of a very selective intervention with high temporal precision, minimizing off target effects typical of pharmacological substances. However, there is already experimental support for the involvement of the OFC-DS pathway in other models of drug-adaptive behaviour. Specifically, acute injection of cocaine activates OFC-DS pathway and induces synaptic potentiation ^34,44^. After short withdrawal, the synaptic transmission between OFC and DS is stronger than in basal condition ^9^. During cue-induced craving, OFC is hyperactive in rodents ^45^ and in human patients with substance use disorders ^46^. These data suggest that the potentiation at OFC-DS pathway is involved in drug seeking behaviour when drug-associated cue is presented. With introduction of a punishment risk, our current data suggest that this OFC-DS hyperfunction is maintained only in compulsive animals. On the other hand, it has also been reported that the OFC is hypoactive in addicted patients ^47^ or in rodents after cocaine self-administration ^48^. However, in these studies, OFC activity was measured only after long withdrawal without any drug related cue presentation, which does not directly contradict a model of gain of function of OFC in addiction prior to drug withdrawal.

In summary, we identified a neural correlate of compulsive reward seeking that is expressed as enhanced synaptic transmission in cortico-striatal synapses and causes a peak in DS calcium activity at the moment of behavioral expression of the integrated reward value. In renouncing mice, the reward-related activity in the DS weakens as punishment arises, likely lowering integrated reward value and thus bringing oDASS to a halt. In compulsive animals on the other hand, the neuronal activity encoding integrated reward value remains intact despite the introduction of punishment and the decision remains in favour of seeking the reward. The identification of OFC-DS neural activity driving pathological decision-making could offer new circuit-based strategies for addiction therapies.

## Acknowledgments

We thank Meaghan Creed for comments on the manuscript; C.Gerfen for providing Cre-mouse lines through the MMRC repository. This work was financed by a grant from the Swiss National Science Foundation (Grant to CL), the National Center of Competence in Research (NCCR) SYNAPSY-The Synaptic Bases of Mental Diseases, and an advanced grant from the European Research Council (MeSSI).

## Author Contributions

MH performed patch recordings, *in vivo* photometry and anatomical tracing with the help of PV. MH and AH did surgeries for viral infection. JF implemented the clustering analysis. CL supervised the work and prepared the manuscript with the help of all authors.

## Author information

1 Dept. of Basic Neurosciences, Faculty of Medicine, University of Geneva, 1 Michel Servet, CH-1211 Geneva, Switzerland.

2 Clinic of Neurology, Dept. of Clinical Neurosciences, Geneva University Hospital, CH-1211 Geneva, Switzerland.

## Data availability

The raw data will be made available under 10.5281/zenodo.4452698

## Extended Data Figures and legends

**Extended Data Fig. 1.**
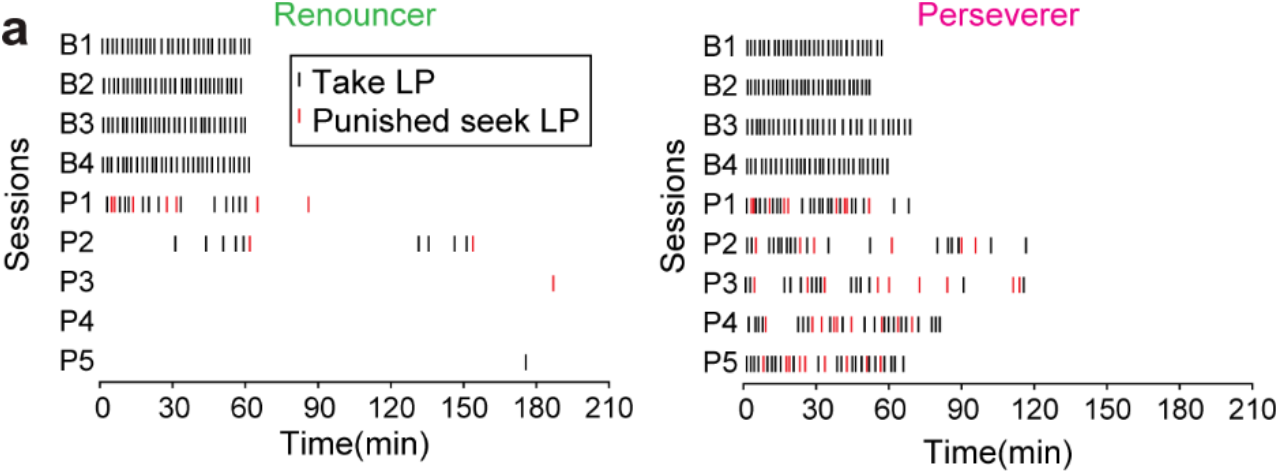
Examples of outcomes of reward seeking behaviour from two mice. **a,** Raster plots showing two mice, one reducing seeking responses facing foot shock (renouncer, left) and one keeping sufficient number of seeking lever presses to complete most of the trials despite negative consequences (perseveres, right). Taking lever presses, followed by the laser stimulation and seeking lever presses triggering foot shock are shown as a function of time for four baseline sessions (B1-B4) and five punishment sessions (P1-P5). Note that seeking lever presses which did not initiate foot shock are not shown.

**Extended Data Fig. 2.**
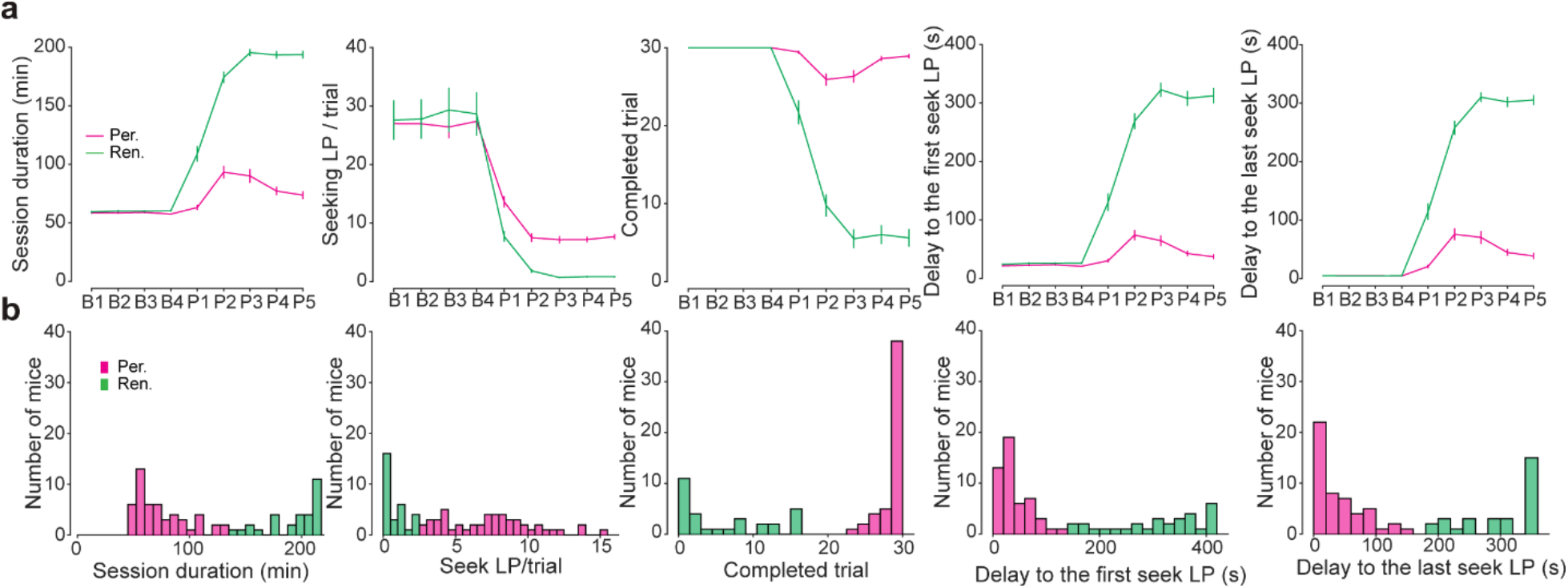
Behavioural parameters of perseveres and renouncers. **a,** Five behavioural parameters during baseline and punishment sessions. **b,** Histograms of behavioural parameters. The average of P4 and P5 data are plotted. Error bars, s.e.m.

**Extended Data Fig. 3.**
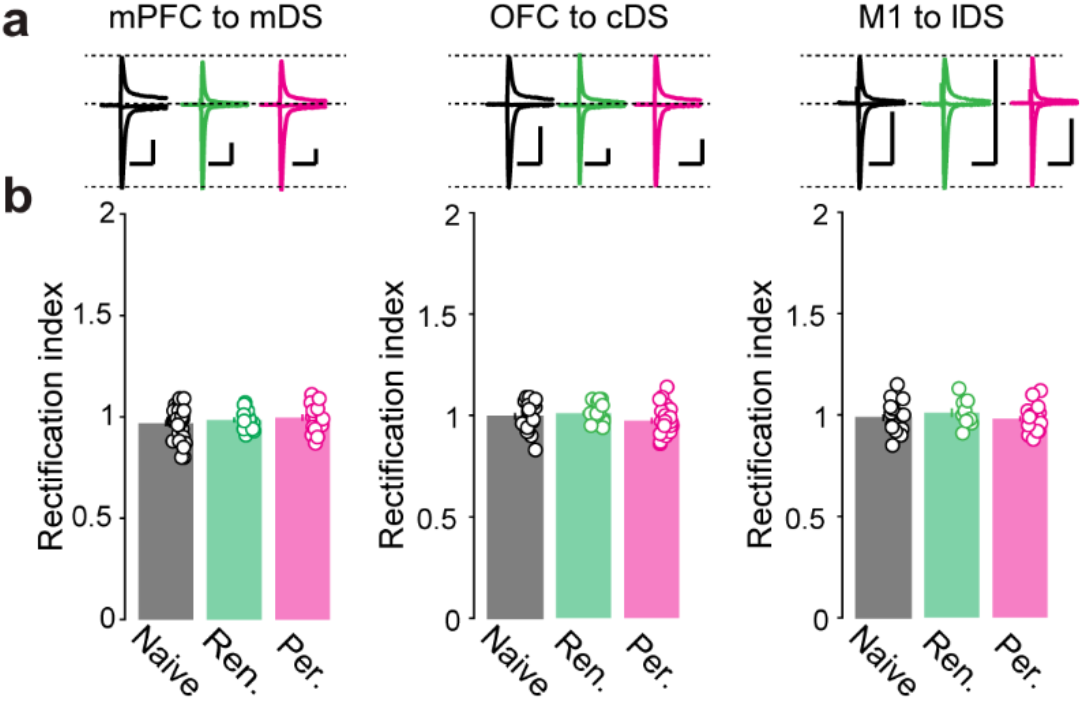
Rectification index at three synapses. **a,** Example traces (Average of 20 sweeps) of AMPAR current recorded at-70,0 and +40 mV. **b,** No significant differences in rectification index were found at any three synapses. mPFC-mDS; F (2, 59) = 0.8481, P=0.4334, Number of recorded neurons; naïve/renounver/perseverer; 31/14/17. OFC-cDS; F (2, 56) = 1.641, P=0.2029, Number of recorded neurons; naïve/renounver/perseverer; 19/17/23. M1-lDS; F (2, 45) = 0.6601, P=0.5217, Number of recorded neurons; naïve/renounver/perseverer; 19/9/20. Scale bars, 50 ms, 500 pA. Error bars, s.e.m.

**Extended Data Fig. 4.**
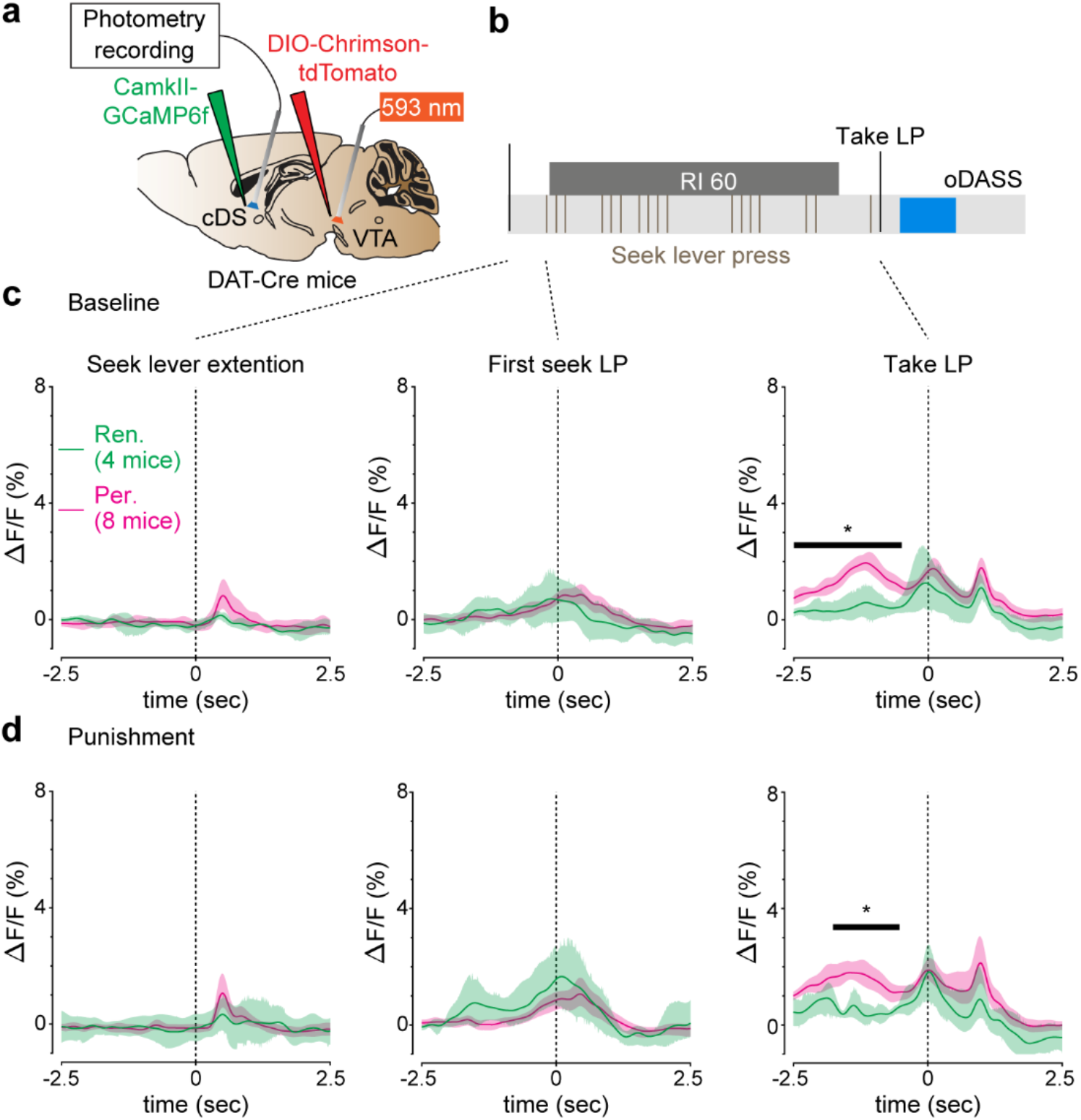
Calcium signals in the cDS during oDASS seeking. **a,** Schematic for the preparation. **b,** Schematic for the oDASS seeking paradigm. **c,** Calcium signals during baseline sessions. Shaded area represents 95% confidence interval. Black line and asterisk indicate time window with statistical difference. Significant difference was detected before take LP, at −2.7089 to −0.4867 s. **d,** Calcium signals during punishment sessions. Shaded area represents 95% confidence interval. Black line and asterisk indicate time window with statistical difference. Significant difference was detected before take LP, at −1.7650 to −0.5261 s.

**Extended Data Fig. 5.**
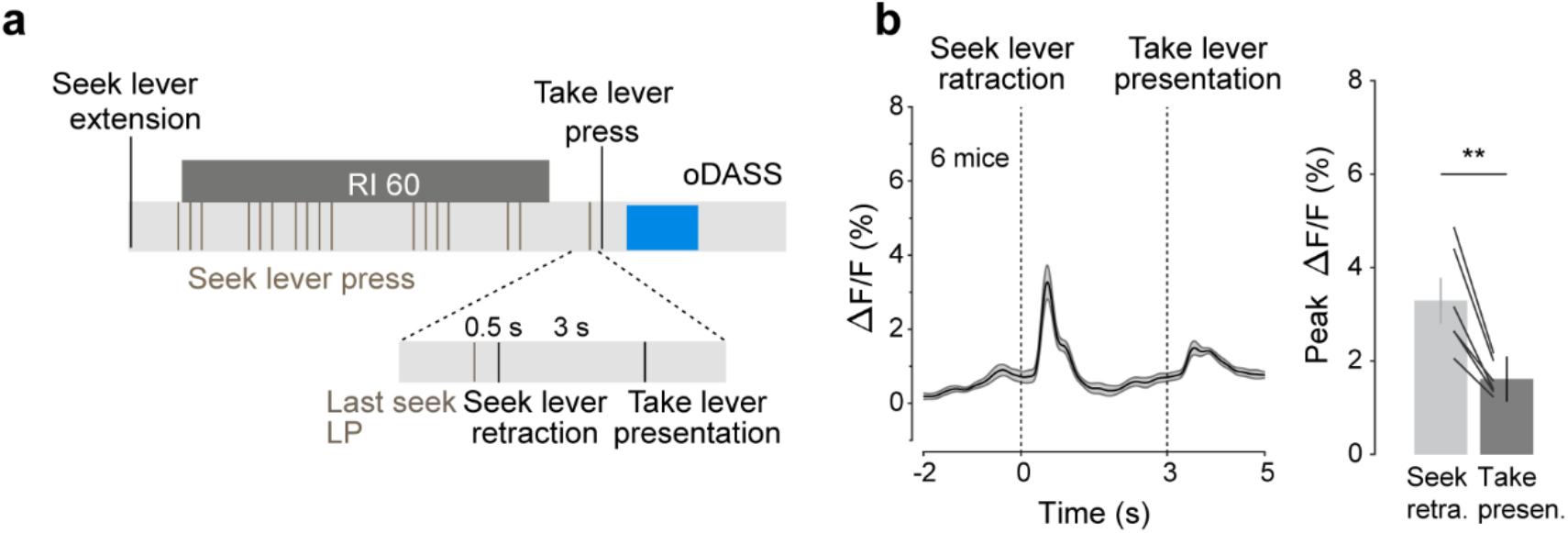
Seek lever retraction, not take lever presentation, induced robust calcium signal. **a,** Schematic for the behavioural paradigm. The delay between the seek lever retraction and the take lever presentation was extended to 3 seconds. **b,** Calcium signals around seek lever retraction and take lever presentation (left). The peak amplitude was higher around the seek lever retraction than around the take lever presentation (t_5_=5.473, P=0.0028). **P<0.01. Error bars, s.e.m.

**Extended Data Fig. 6.**
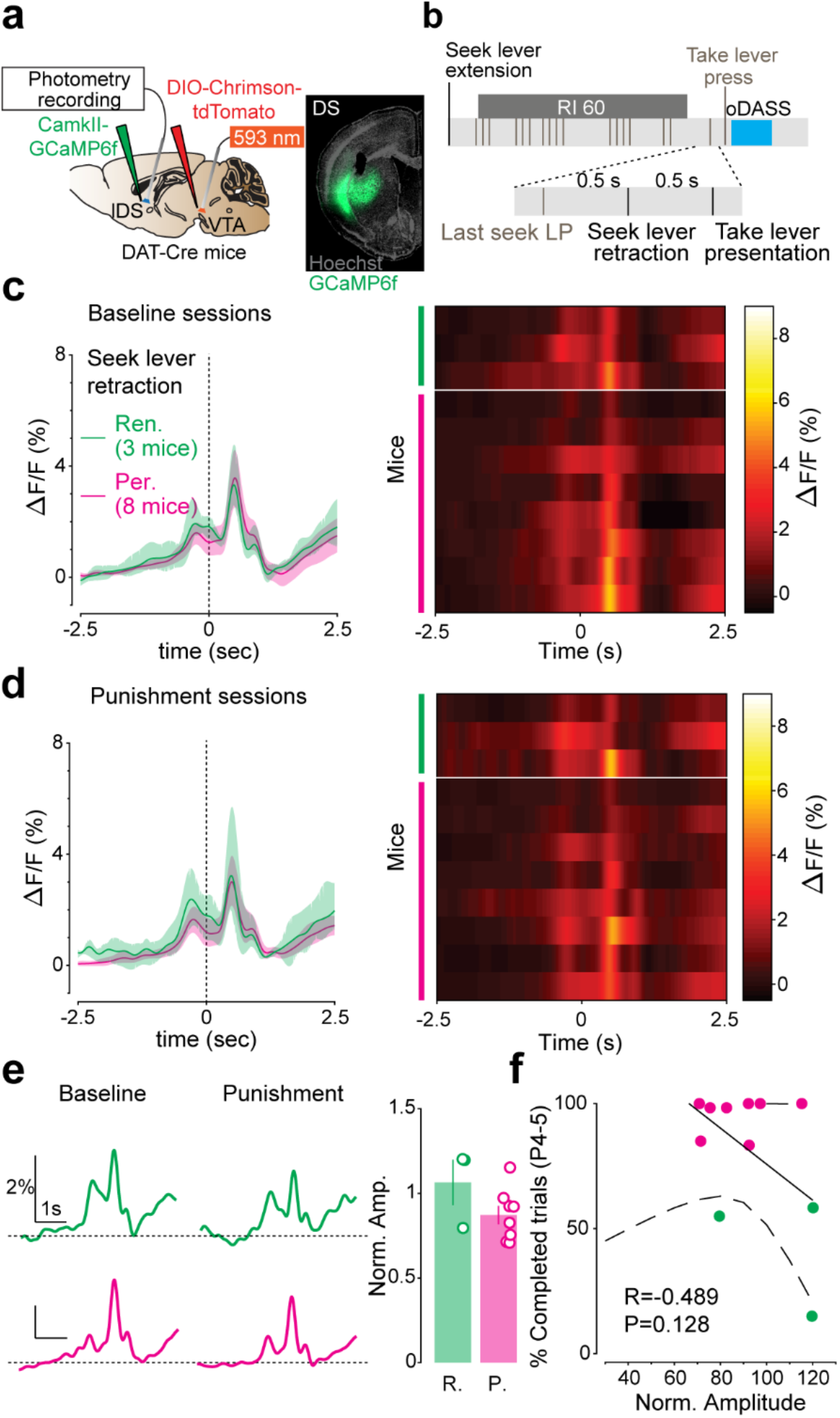
No difference in calcium signals in lDS around seek lever retraction between perseverers and renouncers. **a,** Schematic of the preparation for in vivo photometry recording (left), and coronal image of a mouse brain slice infected with AAV5-CamkII-GCaMP6f in lDS (right). **b,** Schematic for the behavioural paradigm. Calcium signals (ΔF/F) around seek lever retraction are shown in the panels below. **c,** Calcium signal (ΔF/F) around the seek lever retraction during baseline sessions for persevering and renouncing mice (left). Shaded area represents 95% confidence interval. Statistical difference between perseverers and renouncers was not detected. The heatmap shows calcium signal for individual mouse (right). **d,** Calcium signal (ΔF/F) around the seek lever retraction during punishment sessions for persevering and renouncing mice (left). Shaded area represents 95% confidence interval. Statistical difference between perseverers and renouncers was not detected. The heatmap shows calcium signal for individual mouse (right). **e,** Example calcium signals for renouncer (green) and perseverer (pink) around seek lever retraction during baseline and punishment sessions (left). Peak amplitude during punishment sessions is normalized to the amplitude during baseline sessions. The change of peak amplitudes is not stastically significant. (right) (P= 0.1346, t_9_=1.644). **f,** Percent of completed trials (P4-5) as a function of normalized amplitude of calcium signals around seek lever retraction. Percent of completed trials is not correlated to normalized amplitude (Pearson correlation coefficient (R)=−0.489, P=0.128). Error bars, s.e.m.

**Extended Data Fig. 7.**
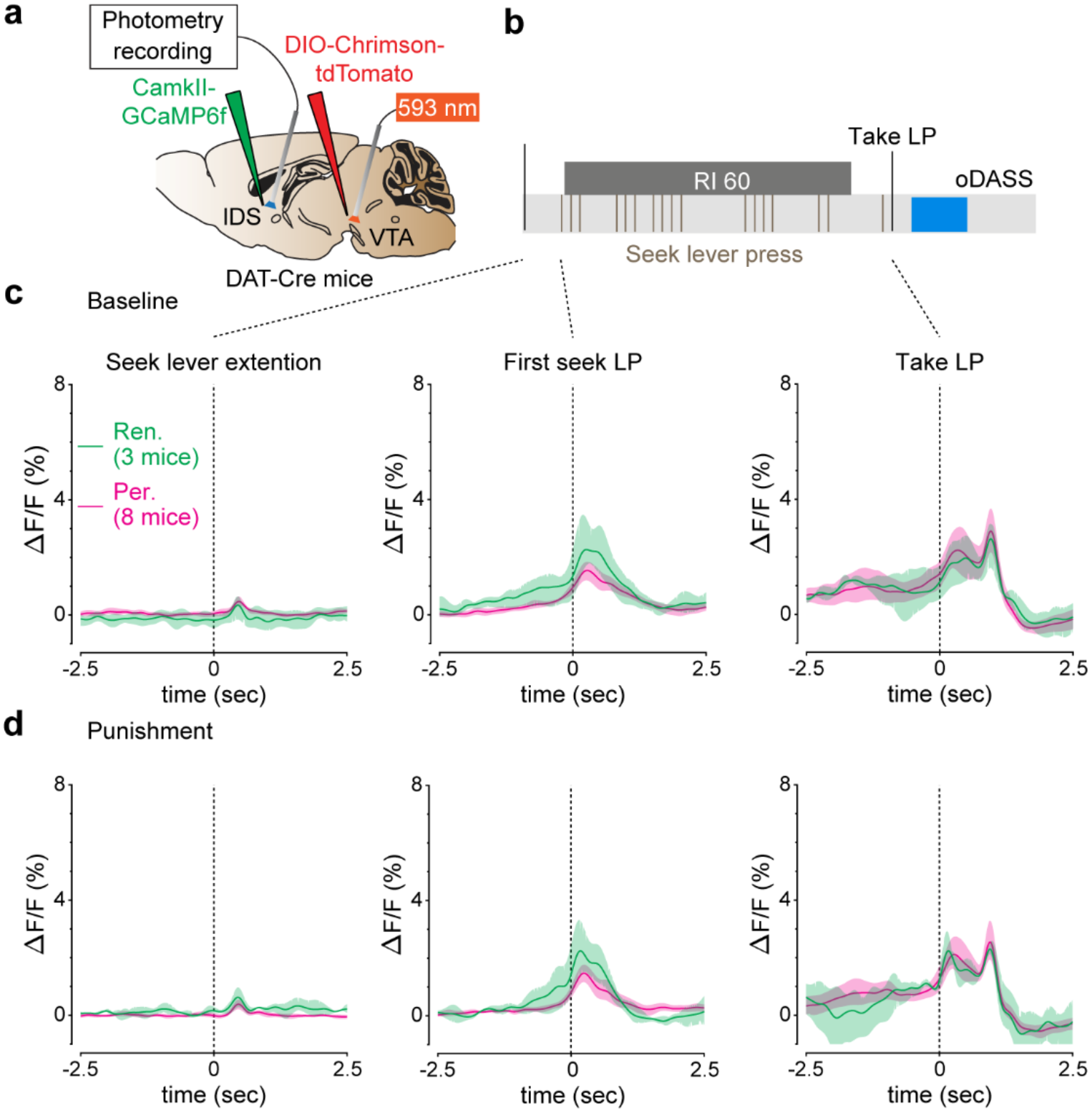
Calcium signals in the lDS during oDASS seeking. **a,** Schematic for the preparation. **b,** Schematic for the oDASS seeking paradigm. **c,** Calcium signals during baseline sessions. Shaded area represents 95% confidence interval. Statistical difference between persevers and renouncers was not detected. **d,** Calcium signals during punishment sessions. Shaded area represents 95% confidence interval. Statistical difference between persevers and renouncers was not detected. Note that calcium signals around the seek lever retraction are show in Extended Data Fig.6.

**Extended Data Fig. 8.**
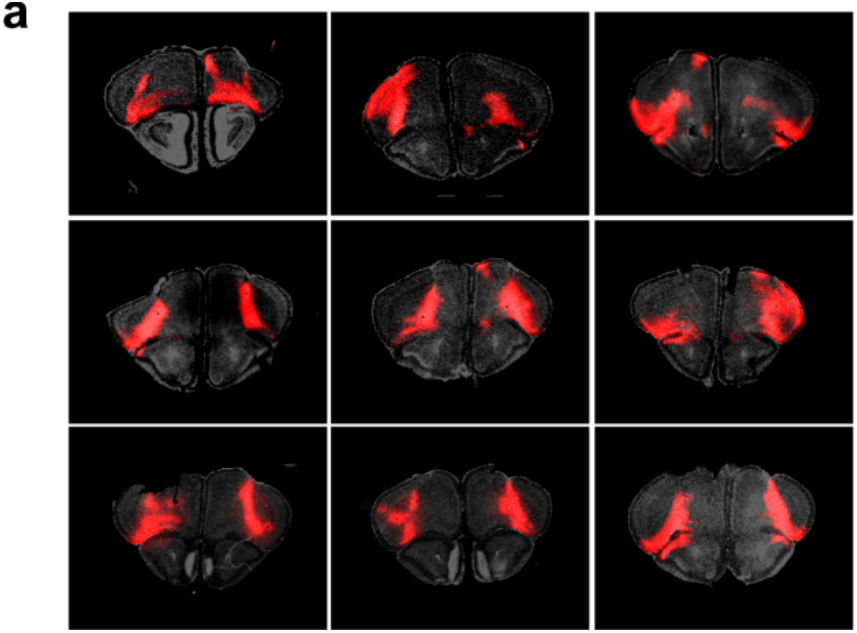
Expression of hM4D in the OFC. **a,** The expression of hM4D in the mice of Fig. 6 c-f. The expression was observed in the orbitofrontal cortex (OFC) of all the mice.

**Extended Data Fig. 9.**
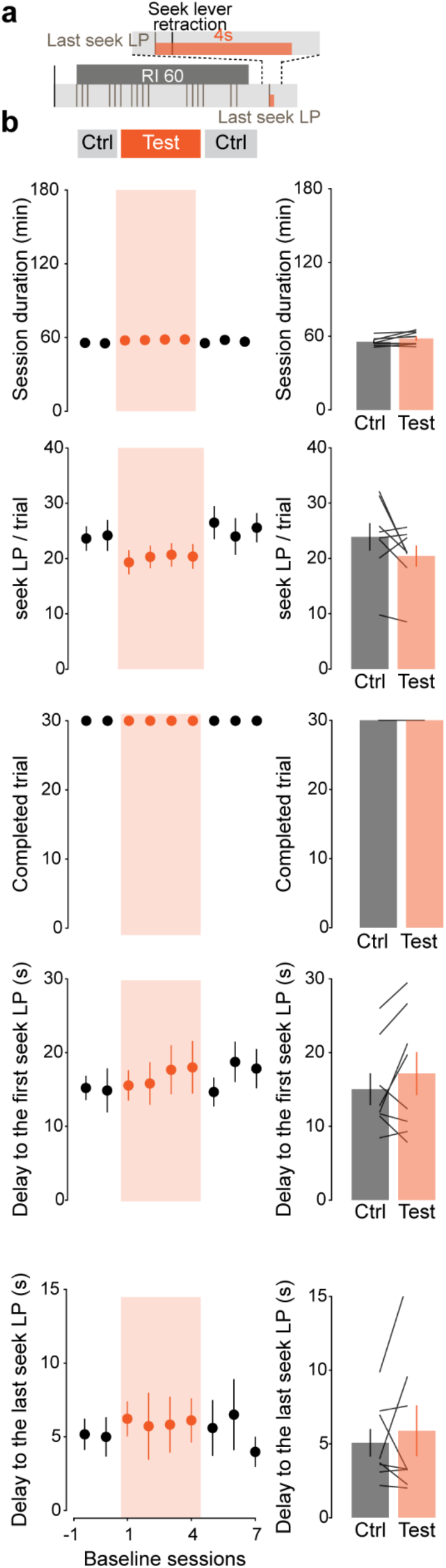
Time locked inhibition in the cDS did not modify reward seeking behavior not associated with punishment. **a,** Schematic for the inhibition protocol. Inhibition was applied at the first seek lever press after the end of RI, as in Fig 7 **d** top. **b,** The same parameters as in Fig. 7 **d,e.**(left). Average of two sessions before inhibition (session −1 and 0) and last three days of inhibition (session 2 to 4) are shown as Ctrl and Test, respectively (bottom, right). Session duration, t_7_=1.775, P=0.1191. seek LP/trial, t_7_=1.807, P= 0.1138. Number of completed trials, zero difference (Wilcoxon test). Delay to the first seek LP, t_7_=1.284, P= 0.2400. Delay to the last seek LP, t_7_=0.6780, P= 0.5196. Error bars, s.e.m.

## Methods

### Animals

Mice (age 8–24 weeks) were heterozygous BAC-transgenic mice in which the Cre-recombinase expression was under the control of the regulatory elements of the dopamine transporter gene (DAT-Cre mice). Dat-Cre mice were originally provided by G. Schutz and only heterozygous mice were used for experiments. DAT B6.SJL-Slc6a3tm1.1(cre)Bkmn/J (also known as DAT-IRES-Cre) mice were also used. Weights and genders were distributed homogeneously among the groups. Transgenic mice had been backcrossed into the C57BL/6 line for a minimum of four generations. All animals were kept in a temperature- and humidity-controlled environment with an inverted 12-h light/12-h dark cycle (lights on at 7PM). All procedures were approved by the Institutional Animal Care and Use Committee of the University of Geneva and by the animal welfare committee of the Cantonal of Geneva, in accordance with Swiss law.

### Stereotaxic injections

AAV5-EF1a-DIO-ChR2(H134R)-eYFP produced at the University of North Carolina (UNC Vector Core Facility) were injected into the VTA. Anaesthesia was induced at 2% and maintained at 1.0% isoflurane (w/v) (Baxter) during surgery. The mouse was placed in a stereotaxic frame (Angle One) and craniotomies were performed using stereotaxic coordinates (for VTA: anterior–posterior (AP) −3.3; medial–lateral (ML) −0.9 with a 10° angle; dorsal– ventral (DV) −4.28). Injections of virus (0.7 μl) used graduated pipettes (Drummond Scientific), broken back to a tip diameter of 10–15 mm, at an infusion rate of 0.05 μl min−1. Following the same procedure, cholera-toxin subunit B (CTB) conjugated with Alexa fluorophore 488 (Thermo Fisher Scientific, Massachussetts, USA), 555 or 647 was unilaterally (left/right hemisphere, counterbalanced) injected in the medial (AP +0.7; ML ±1.1; DV −2.9), central (AP +0.8; ML ±1.9; DV −3.4) or lateral (AP +0.7; ML ±2.5; DV −2.1) part of the DS, respectively. AAV8-hSyn-ChrimsonR-tdTomato (UNC) was injected bilaterally in the mPFC (AP +1.9; ML ±0.3; DV −2.5), OFC (AP +2.6; ML ±1.75; DV −1.95) or M1 (AP +1.1; ML ±1.7; DV −1.75). AAV5-CamKII-GCamp6f produced at University of Pennsylvania (UPENN) was unilaterally injected in the cDS (AP +0.8; ML +1.65; DV −3.3). AAV1-CamKII-hM4D-mcherry (UNC) was bilaterally injected in the OFC (AP +2.6; ML ±1.75; DV −1.95). AAV5-CamKII-eArch3.0T-eYFP (UNC) was bilaterally injected in the cDS (AP +0.8; ML +1.65; DV −3.3). During the same surgical procedure, a unique chronically indwelling optic fibre cannula was implanted above the VTA using the exact same coordinates as for the injection except for DV coordinate, which was reduced to 4.18. A photometry fibre (Doric lenses, MFC_400/430-0.48_4mm_ZF2.5(G)_FLT) was implanted unilaterally in the cDS (AP +0.8; ML 1.65; DV −3.1). Optic fibres for eArchT3.0 stimulation were placed in each cDS (AP +0.8; ML ±1.65; DV −3.1). Two screws were fixed into the skull to support the implant, which was further secured with dental cement. The first behavioural session typically occurred 10−14 days after surgery to allow sufficient expression of ChR2.

### Optogenetic self-stimulation apparatus

Mice infected with AAV5-EF1a-DIO-ChR2(H134R)-eYFP or AAV8-hSyn-Flex-ChrimsonR-tdTomato in the VTA were placed during their dark phase in operant chambers (ENV-307A-CT, Med Associates) situated in a sound-attenuating box (Med Associates). The optic fibre of the mouse was connected to DPSS blue- or orange-light lasers (CNI-473-140-10-LED-TTL1-MM200FC; CNI-593-200-10-LED-TTL1-MM200FC; Laser 2000) via a FC/PC fibre cable (M72L02; Thorlabs) and a simple rotary joint (FRJ-FC-FC; Doric Lenses) allowing free movement during operant behaviour. Power at the exit of the patch cord was set to 15 ± 1 mW. Two retractable levers were present on one wall of the chamber and a cue light was located above each lever. A rack mount interface cabinet (SG-6010A, Med Associates) containing a programmable constant current shocker (ENV-413, Med Associates) connected to a quick disconnect grid harness (ENV-307A-QD, Med Associates) was used to provide foot shocks during punishment sessions. The apparatus was controlled, and data captured using MED-PC IV software (Med Associates). For acute inhibition in the cDS, a double rotary joint (FRJ_1×2i_FC-2FC, Doric Lenses) was used to connect cables to each hemisphere. For orange laser stimulation of eArchT3.0 or ChrimsonR, a shutter (CMSA-SR475_FC, Doric Lenses) was used to avoid variation in intensities during laser warm-up.

### Acquisition of taking response

In this first phase of training, each session started with the presentation of the take lever (left/right, counterbalanced), which then is never retracted during the session. A single press on the take lever (FR1) resulted in a 10-s illumination of a cue light (pulses of 1 s at 1 Hz). After a delay of 5 s, onset of a 15-s laser stimulation composed of 30 bursts separated by 250 ms (each burst consisted of 5 laser pulses of 4-ms pulse width at 20 Hz). A 20-s time-out followed the rewarded lever press, during which lever presses had no consequences but were still recorded. Each of the 5 daily acquisition sessions lasted 120 min or until the mouse reached 80 optogenetic stimulations, whichever came first.

### Training of the seek-take chain

During this second phase, mice learned the seek-take chain with FR1. Each trial of the seek-take chained schedule began with the presentation of the seek lever (opposite to the take lever), with the take lever retracted. A single press on the seek lever resulted in the retraction of the seek lever (500ms later) and presentation of the take lever (again 500ms later). A single press on the take lever resulted in the delivery of the cue light immediately after the press, retraction of the take lever (500 ms later) and laser stimulation (5s later) that last 15s. After a 5s time-out, another trial started the extension of the seek lever. Daily session lasted 120 min or until the mouse reached 80 optogenetic stimulations, whichever came first. Mice underwent 7 sessions of FR1 seek-take chained schedule, and then a random interval (RI) schedule was introduced.

During the RI schedule, each trial started with the presentation of the seek lever, with the take lever retracted. The first press on the seek lever initiated the RI schedule. Seek lever presses during RI had no programmed consequences but were counted. The first seek lever press after the end of the RI resulted in the retraction of the seek lever and the presentation of the take lever. A single press on the take lever resulted in the delivery of the cue light, laser stimulation and the retraction of the take lever. Following a 5s time-out, the seek lever presentation initiated another trial. Three RI parameters were used: RI5 (0.5s, 5s or 10s), RI30 (15s, 30s or 45s) and RI60 (45s, 60s or 75s). Mice underwent 3 sessions for each RI. Sessions lasted 360 min or until the mouse reached 80, 60 or 40 optogenetic stimulations, whichever came first, for respectively RI5, 30 or 60.

### Evaluation of compulsivity

After the training, mice underwent additional sessions of RI60 schedule. Stable performance (earning all 30 stimulations) across four consecutive sessions served as a baseline before the punishment sessions. During punishment sessions, for 30% of the trials, the first seek lever press after RI resulted in the retraction of the seek lever and the delivery of the mild foot shock (0.25mA, 500ms) and led to the time-out period directly without the presentation of the take lever. The remaining 70% trials contained no punishment but resulted in the extension of the take lever, press on which resulted in the delivery of the laser stimulation. Mice were given 5 punishment sessions. During baseline and punishment sessions, mice had a maximum of 7min, including the RI60 duration to complete each trial. After a failed trial (no seek press to start a RI after seek lever extension or no seek press after the end of the RI in the allowed 7 min), a novel seek lever presentation is given.

### Slice electrophysiology

Coronal 230-μm slices of the DS were prepared in cooled artificial cerebrospinal fluid containing (in mM): NaCl 119, KCl 2.5, MgCl 1.3, CaCl2 2.5, Na2HPO4 1.0, NaHCO3 26.2 and glucose 11, bubbled with 95% O2 and 5% CO2. Slices were kept in ACSF at 35℃ for 15-30 min for recovery and then kept at room temperature until recordings. Once transferred in the recording chamber, slices were superfused with 2.5 ml/min artificial cerebrospinal fluid at 32–34℃.

#### Ex vivo synaptic properties of the striatum

Visualized whole-cell patch-clamp recording techniques were used to measure synaptic responses to optogenetic stimulation of mPFC,OFC or M1 terminals infected with ChrimsonR. The access resistance was monitored by a hyperpolarizing step of −14 mV. The liquid junction potential was small (−3 mV), and therefore traces were not corrected. Experiments were discarded if the access resistance varied by more than 20%. Currents were amplified (Multiclamp 700B, Axon Instruments), filtered at 5 kHz and digitized at 20 kHz (National Instruments Board PCI-MIO-16E4, Igor, Wave Metrics). For recordings of optogenetically evoked EPSCs, the internal solution contained (in mM): CsCl 130, NaCl 4, creatine phosphate 5, MgCl2 2, NA2ATP 2, NA3GTP 0.6, EGTA 1.1, HEPES 5 and spermine 0.1. QX-314 (5 mM) was added to the solution to prevent action currents. Synaptic currents were evoked by short light pulses (4 ms) at 0.1 Hz through an LED (PE-100, cool-LED; 585nm) placed through the objective above the tissue. The holding potential was at +40mV to measure the AMPAR/NMDAR ratio. To isolate AMPAR-evoked EPSCs the NMDA antagonist d-2-amino-5-phosphonovaleric acid (D-AP5, 50 μM) was applied to the bath. The NMDAR component was calculated as the difference between the EPSCs measured in the absence and in the presence of D-AP5. The AMPAR/NMDAR ratio was calculated by dividing the peak amplitudes. The rectification index of AMPAR was calculated as the ratio of the chord conductance calculated at negative potential (−70mV) divided by chord conductance at positive potential (+40mV). Examples traces are averages of 10–15 sweeps. All experiments were performed in the presence of picrotoxin (100 μM).

#### Functional connectivity

For recording EPSCs from mPFC, OFC or M1 to striatal MSNs, the internal solution contained; (in mM): CsCl 130, NaCl 4, creatine phosphate 5, MgCl2 2, NA2ATP 2, NA3GTP 0.6, EGTA 1.1, HEPES 5 and spermine 0.1, QX-314 5. Picrotoxin (100 μM) was added to the bath perfusion to block inhibitory currents during the recordings. The holding potential was at – 70mV. Synaptic currents were evoked by 2 short light pulses (4ms duration, 76ms interval) at 0.1Hz. The amplitude of the first pulse is shown in Figure 3.

#### DREADD validation

For recording EPSCs from OFC to striatal MSNs, the internal solution contained; (in mM): CsCl 130, NaCl 4, creatine phosphate 5, MgCl2 2, NA2ATP 2, NA3GTP 0.6, EGTA 1.1, HEPES 5 and spermine 0.1, QX-314 5. Picrotoxin (100 μM) was added to the bath perfusion to block inhibitory currents during the recordings. The holding potential was at −70mV. Synaptic currents were evoked by 2 short light pulses (4ms duration, 76ms interval) at 0.1Hz. After obtaining 5 min stable baseline, CNO (CNO, 10 μM) was applied to the bath. The magnitude of reduction of EPSCs was determined by comparing the average EPSCs that were recorded 15-20 min after CNO application to average EPSCs recorded immediately before CNO application.

#### eArchT3.0 validation

For recordings of striatal MSNs infected with eArchT3.0 the internal solution contained (in mM): potassium gluconate 130, MgCl2 4, Na2ATP 3.4, Na3GTP 0.1, creatine phosphate 10, HEPES 5 and EGTA 1.1. Firing was triggered by a current step (5s duration, 350pA step) with or without stimulation of eArchT3.0 (4s) with the LED (PE-100, cool-LED;585nm). Picrotoxin (100 μM) and CNQX (10 μM) were added to the bath perfusion to block excitatory and inhibitory synaptic transmission, respectively.

### Fibre photometry recordings

Fibre photometry recording were performed during baseline and punishment sessions. Dat-Cre mice infected with AAV8-hSyn-Flex-ChrimsonR-tdTomato (UNC) in the VTA and with AAV5-CamkII-GCamp6f (UPENN) in the dorsal striatum were used for the recordings. For DREADD experiments, AAV1-CamKII-hM4D-mcherry (UNC) was also injected in the OFC. CNO (2mg/kg in saline solution) was injected 1 hour prior to each session. Mice were recorded for 30 min maximum per session to minimise bleaching. Striatum neurons were illuminated with blue (470 nm wavelength, M470F3, Thorlabs) and violet (405 nm wavelength, M405FP1, Thorlabs) filtered excitation LED lights, that were sinusoidally modulated at 211 and 531 Hz. Green emission light (500–550 nm) was collected through the same fibre that was used for excitation and passed onto a photo-receiver (Newport 2151, Doric Lenses). Pre-amplified signals were then demodulated by a real-time signal processor (RZ5P, Tucker Davis Systems) to determine contributions from 470 nm and 405 nm excitation sources. TTL signals of the relevant stimuli were directly sent from the operant chamber to the signal processor. Analysis was performed offline in MATLAB. To calculate ΔF/F0, a linear fit was applied to the 405-nm control signal to align it to the 470-nm signal. This fitted 405-nm signal was used as F0 in standard ΔF/F0 normalization ((F(t) − F0(t))/F0(t)). Averaged peristimulus activity traces were then constructed for which the mean baseline fluorescence (5 to 3 s before the relevant event) was subtracted from the trace.

### Acute inhibition of the DS

Dat-Cre mice were infected with AVV5-CamKII-eArchT3.0-eYFP in the cDS and optogenetic fibres were inmpanted bilaterally in the cDS. Orange light laser started at specific epochs of behaviour; (1) Immediately after the first seek press after the end of random interval. Laser was turned on continuously for 4s. (2) Immediately after the first seek press after the presentation of seek lever. Laser was turned on continuously for 4s.

### Tissue preparation for imaging

Mice were anaesthetized with pentobarbital (300 mg/kg, intraperitoneally, Sanofi-Aventis) and transcardially perfused with 4% (w/v) paraformaldehyde in PBS (pH 7.5). Brains were post-fixed overnight in the same solution and stored at 4 °C. Coronal sections (50-mm thick) were cut with a vibratome (Leica), stained with Hoechst (Sigma- Aldrich) and mounted with Mowiol (Sigma-Aldrich). Full images of brain slices were obtained with a Zeiss Axioscan Z1 system equipped with a Plan-Apochromat 10×/0.45 NA objective, together with filters for 4′,6-diamidino-2-phenylindole (DAPI) (emission band-pass filter: 445/50 nm), enhanced green fluorescent protein (eGFP) (emission band-pass filter: 525/50 nm) and cyanine 3 (Cy3) (emission band- pass filter: 605/70 nm).

Images from VTA and striatum were obtained using sequential laser scanning confocal microscopy (Zeiss LSM800). Photomicrographs were obtained with the following band-pass and long-pass filter settings: UV excitation (band-pass filter: 365/12 nm), GFP (band-pass filter: 450−490 nm), Cy3 (band-pass filter: 546/12 nm) and Cy5 (band-pass filter: 546/12 nm). For biocytin (Sigma-Aldrich, B4261) staining, streptavidin–Cy5 (Invitrogen 434316) was used. For immunohistochemistry, the following primary antibody (rabbit polyclonal anti-tyrosine hydroxylase, Millipore AB152, lot 2722866, diluted 1:500) and the secondary antibody (donkey anti-rabbit Cy3, Millipore AP182C, lot 2397069, diluted 1:500) were used.

### Statistics

Statistical analysis was performed in GraphPad Prism 8. For all tests, the threshold of statistical significance was placed at 0.05. For experiments involving two independent subjects or the same subjects at two different time points, two tailed Student’s unpaired or paired t-test was used, respectively. For experiments involving more than two groups, mixed-factor ANOVA defining both between-(for example, inhibition at the last seek LP vs inhibition at the first seek LP) and/or within-(for example, without or with ArchT3.0 inhibition) group factors. Where significant main effects or interaction terms were found (p<0.05), further comparisons were made by a two-tailed Student’s t-test with Bonferroni correction. To compare the number of completed trials (Figure 6E and 7F), Wilcoxon test was used because of its distribution. When multiple comparison was performed, P value was corrected with Bonferroni-Dunn method (Figure 7F). To measure Pearson correlation coefficient and its 95% confidence interval, MATLAB2018b (MathWorks) was used. To compare the photometry signal (bin size=0.0197) between perseverers and renouncers mice at a given event (e.g. at the seek lever retraction), we first computed for each mouse the average signal over trials from −5 to 5 seconds around the event. The obtained average signal of each mouse was then normalized by subtracting the mean over the baseline ranging from −5 to −4 seconds. This produced two matrices, one for perseverers and the other for renouncers for a given event. For each matrix, we computed the 95% bootstrap confidence interval (CI) with 10’000 sample with replacement (Matlab function ‘bootci’; bias corrected percentile method, alpha=0.05). For a given event, time periods longer than 0.6 seconds for which the CI of perseverers and renouncers continuously do not overlap are consider as significant.

### Identification of perseverers and renouncers mice

In order to identify perseverers and renouncers mice, we performed a clustering. To this end, five behavioural parameters were used: session duration, number of seeking lever presses, number of completed trials, delay to the first seeking lever press after the extension of the seeking lever and delay to the last seeking lever press after the end of random interval. The number of seeking lever presses were rescaled with a symmetrized log-scale scale 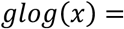 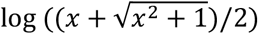 to deal with their large spreading. On the other hand, a standard z-score was used for the other four parameters. We then applied a hierarchical clustering method (Matlab functions ‘pdist’, ‘linkage’, ‘cluster’ with a squared euclidean metric and ward linkage distance) then set a cut-off to the tree to have two clusters (the renouncer and perseverer mice). The degree of robustness of the clustering was assessed by computing the mean silhouette score across clusters. To check if the clustering obtained is significantly different from chance, we compared our result to a data distribution obtained by shuffling the mice indices for each variable 1000x, redo the clustering and compute the robustness for each sample.

---

**Table 1.**
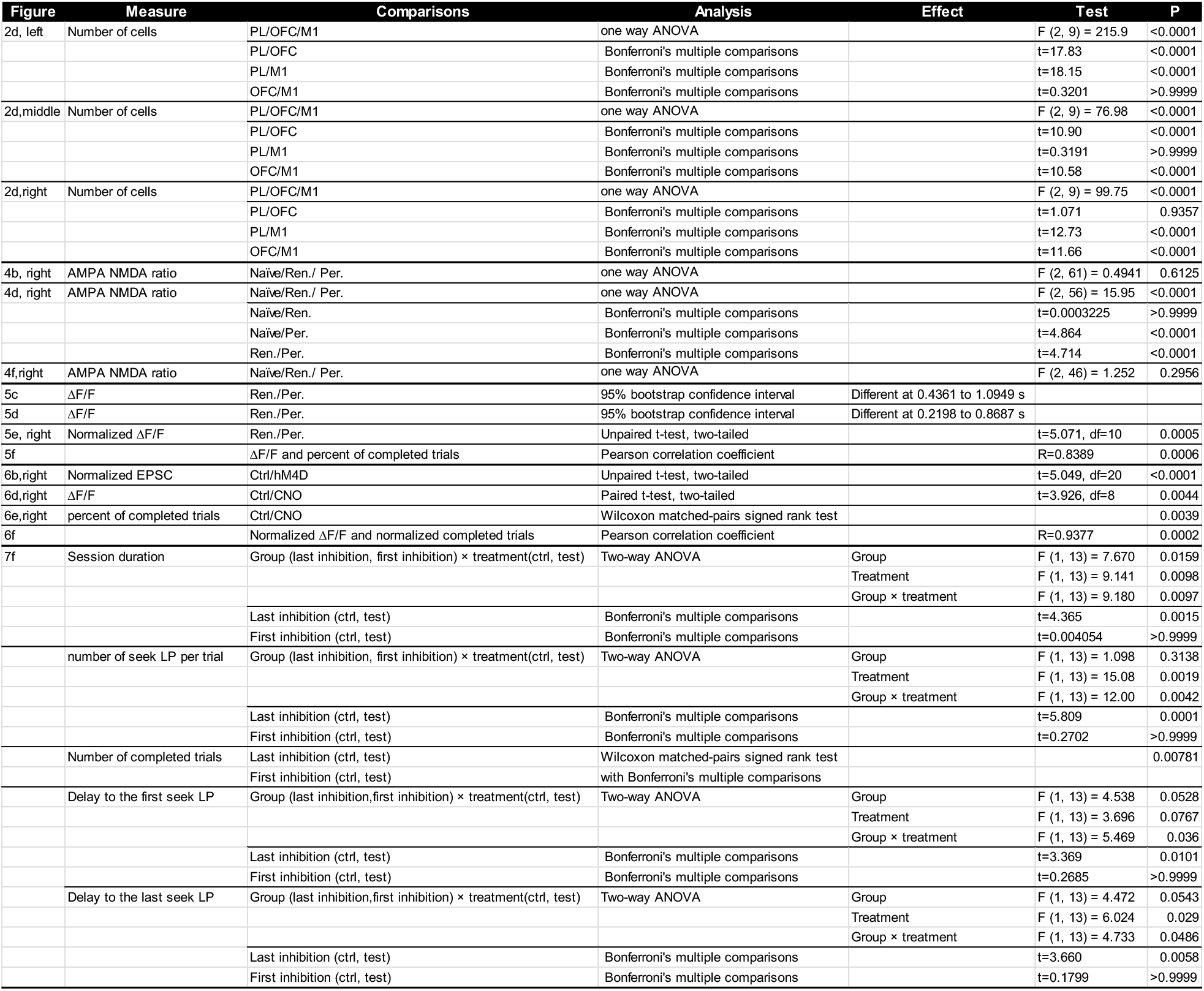
Main Figures Statistic Table.

**Table 2.** Extended Data Figures Statistic Table.

| Figure | Measure | Comparisons | Analysis | Effect | Test | P |
| --- | --- | --- | --- | --- | --- | --- |
| Ex. Fig. 3b, left | Rectification index | Naïve/Ren./ Per. | one way ANOVA |  | F (2, 59) = 0.8481 | 0.4334 |
| Ex. Fig. 3b, middle | Rectification index | Naïve/Ren./ Per. | one way ANOVA |  | F (2, 56) = 1.641 | 0.2029 |
| Ex. Fig. 3b, right | Rectification index | Naïve/Ren./ Per. | one way ANOVA |  | F (2, 45) = 0.6601 | 0.5217 |
| Ex. Fig. 4b, top, left | $\Delta F/F$ | Ren./Per. | 95% bootstrap confidence interval | Difference was not detected | | |
| Ex. Fig. 4b, top, middle | $\Delta F/F$ | Ren./Per. | 95% bootstrap confidence interval | Difference was not detected | | |
| Ex. Fig. 4b, top, right | $\Delta F/F$ | Ren./Per. | 95% bootstrap confidence interval | Different at -2.7089 to -0.4867 | | |
| Ex. Fig. 4b, bottom, left | $\Delta F/F$ | Ren./Per. | 95% bootstrap confidence interval | Difference was not detected | | |
| Ex. Fig. 4b, bottom, middle | $\Delta F/F$ | Ren./Per. | 95% bootstrap confidence interval | Difference was not detected | | |
| Ex. Fig. 4b, bottom, right | $\Delta F/F$ | Ren./Per. | 95% bootstrap confidence interval | Different at -1.7650 to -0.5261 | | |
| Ex. Fig. 5b, right | $\Delta F/F$ | seek ret./take pre. | paired t-test | | t=5.473, df=5 | 0.0028 |
| Ex. Fig. 6c | $\Delta F/F$ | Ren./Per. | 95% bootstrap confidence interval | Difference was not detected | | |
| Ex. Fig. 6d | $\Delta F/F$ | Ren./Per. | 95% bootstrap confidence interval | Difference was not detected | | |
| Ex. Fig. 6e, right | Normalized $\Delta F/F$ | Ren./Per. | Unpaired t-test, two-tailed | | t=1.644, df=9 | 0.1346 |
| Ex. Fig. 6f | | $\Delta F/F$ and percent of completed trials | Pearson correlation coefficient | | R=-0.4892 | 0.1267 |
| Ex. Fig. 7b, top, left | $\Delta F/F$ | Ren./Per. | 95% bootstrap confidence interval | Difference was not detected | | |
| Ex. Fig. 7b, top, middle | $\Delta F/F$ | Ren./Per. | 95% bootstrap confidence interval | Difference was not detected | | |
| Ex. Fig. 7b, top, right | $\Delta F/F$ | Ren./Per. | 95% bootstrap confidence interval | Difference was not detected | | |
| Ex. Fig. 7b, bottom, left | $\Delta F/F$ | Ren./Per. | 95% bootstrap confidence interval | Difference was not detected | | |
| Ex. Fig. 7b, bottom, middle | $\Delta F/F$ | Ren./Per. | 95% bootstrap confidence interval | Difference was not detected | | |
| Ex. Fig. 7b, bottom, right | $\Delta F/F$ | Ren./Per. | 95% bootstrap confidence interval | Difference was not detected | | |
| Ex. Fig. 9b | Session duration | ctrl/test | paired t-test |  | t=1.775, df=7 | 0.1191 |
|  | number of seek LP per trial | ctrl/test | paired t-test |  | t=1.807, df=7 | 0.1138 |
|  | Number of completed trials | ctrl/test | Wilcoxon test |  | zero difference |  |
|  | Delay to the first seek LP | ctrl/test | paired t-test |  | t=1.284, df=7 | 0.24 |
|  | Delay to the last seek LP | ctrl/test | paired t-test |  | t=0.6780, df=7 | 0.5196 |

## References

1. Lüscher, C., Robbins, T. W. & Everitt, B. J. The transition to compulsion in addiction. Nat. Rev. Neurosci. 21, 247–263 (2020).

2. Anthony, J. C., Warner, L. A. & Kessler, R. C. Comparative Epidemiology of Dependence on Tobacco, Alcohol, Controlled Substances, and Inhalants: Basic Findings From the National Comorbidity Survey. Exp. Clin. Psychopharmacol. 2, 244–268 (1994).

3. Pelloux, Y., Dilleen, R., Economidou, D., Theobald, D. & Everitt, B. J. Reduced forebrain serotonin transmission is causally involved in the development of compulsive cocaine seeking in rats. Neuropsychopharmacology 37, 2505–2514 (2012).

4. Deroche-Gamonet, V., Belin, D. & Piazza, P. V. Evidence for addiction-like behavior in the rat. Science 305, 1014–1017 (2004).

5. Pascoli, V., Terrier, J., Hiver, A. & Lüscher, C. Sufficiency of Mesolimbic Dopamine Neuron Stimulation for the Progression to Addiction. 88, Neuron 1054–1066 (2015).

6. Pelloux, Y., Murray, J. E. & Everitt, B. J. Differential vulnerability to the punishment of cocaine related behaviours: Effects of locus of punishment, cocaine taking history and alternative reinforcer availability. Psychopharmacology (Berl). 232, 125–134 (2015).

7. Siciliano, C. A. et al. A cortical-brainstem circuit predicts and governs compulsive alcohol drinking. Science 366, 1008–1012 (2019).

8. Kasanetz, F. et al. Transition to addiction is associated with a persistent impairment in synaptic plasticity. Science 328, 1709–1712 (2010).

9. Hu, Y. et al. Compulsive drug use is associated with imbalance of orbitofrontal- and prelimbic-striatal circuits in punishment-resistant individuals. Proc. Natl. Acad. Sci. 116, 9066–9071 (2019).

10. Everitt, B. J. & Robbins, T. W. Drug Addiction: Updating Actions to Habits to Compulsions Ten Years On. Annu. Rev. Psychol. 67, 23–50 (2016).

11. Lüscher, C. & Ungless, M. A. The mechanistic classification of addictive drugs. PLoS Med. 3, 2005–2010 (2006).

12. Di Chiara, G. & Imperato, A. Drugs abused by humans preferentially increase synaptic dopamine concentrations in the mesolimbic system of freely moving rats. Proc. Natl. Acad. Sci. U. S. A. 85, 5274–5278 (1988).

13. Bocklisch, C. et al. Cocaine disinhibits dopamine neurons by potentiation of GABA transmission in the ventral tegmental area. Science 341, 1521–1525 (2013).

14. Pascoli, V. et al. Contrasting forms of cocaine-evoked plasticity control components of relapse. Nature 509, 459–464 (2014).

15. Pascoli, V., Turiault, M. & Lüscher, C. Reversal of cocaine-evoked synaptic potentiation resets drug-induced adaptive behaviour. Nature 481, 71–75 (2012).

16. Ma, Y. et al. Bidirectional Modulation of Incubation of Cocaine Craving by Silent Synapse-Based Remodeling of Prefrontal Cortex to Accumbens Projections. Neuron 83, 1453–1467 (2014).

17. Hearing, M. C. et al. Reversal of morphine-induced cell-type–specific synaptic plasticity in the nucleus accumbens shell blocks reinstatement. Proc. Natl. Acad. Sci. U. S. A. 113, 757–762 (2016).

18. Willuhn, I., Burgeno, L. M., Everitt, B. J. & Phillips, P. E. M. Hierarchical recruitment of phasic dopamine signaling in the striatum during the progression of cocaine use. Proc. Natl. Acad. Sci. U. S. A. 109, 20703–20708 (2012).

19. Everitt, B. J. & Robbins, T. W. From the ventral to the dorsal striatum: Devolving views of their roles in drug addiction. Neurosci. Biobehav. Rev. 37, 1946–1954 (2013).

20. Zapata, A., Minney, V. L. & Shippenberg, T. S. Shift from Goal-Directed to Habitual Cocaine Seeking after Prolonged Experience in Rats. J. Neurosci. 30, 15457–15463 (2010).

21. Jonkman, S., Pelloux, Y. & Everitt, B. J. Differential Roles of the Dorsolateral and Midlateral Striatum in Punished Cocaine Seeking. J. Neurosci. 32, 4645–4650 (2012).

22. Balleine, B. W. & O’Doherty, J. P. Human and rodent homologies in action control: Corticostriatal determinants of goal-directed and habitual action. Neuropsychopharmacology 35, 48–69 (2010).

23. Belin, D., Jonkman, S., Dickinson, A., Robbins, T. W. & Everitt, B. J. Parallel and interactive learning processes within the basal ganglia: Relevance for the understanding of addiction. Behav. Brain Res. 199, 89–102 (2009).

24. Gruber, A. J. & McDonald, R. J. Context, emotion, and the strategic pursuit of goals: Interactions among multiple brain systems controlling motivated behavior. Front. Behav. Neurosci. 6, 50 (2012).

25. Thorn, C. A., Atallah, H., Howe, M. & Graybiel, A. M. Differential Dynamics of Activity Changes in Dorsolateral and Dorsomedial Striatal Loops during Learning. Neuron 66, 781–795 (2010).

26. Yin, H. H. & Knowlton, B. J. The role of the basal ganglia in habit formation. Nat. Rev. Neurosci. 7, 464–476 (2006).

27. Yin, H. H., Ostlund, S. B., Knowlton, B. J. & Balleine, B. W. The role of the dorsomedial striatum in instrumental conditioning. Eur. J. Neurosci. 22, 513–523 (2005).

28. Hunnicutt, B. J. et al. A comprehensive excitatory input map of the striatum reveals novel functional organization. eLife 5, e19103 (2016).

29. Pascoli, V. et al. Stochastic synaptic plasticity underlying compulsion in a model of addiction. Nature 564, 366–371 (2018).

30. Chen, B. T. et al. Rescuing cocaine-induced prefrontal cortex hypoactivity prevents compulsive cocaine seeking. Nature 496, 359–362 (2013).

31. Domingo-rodriguez, L. et al. A specific prelimbic-nucleus accumbens pathway controls resilience versus vulnerability to food addiction. Nat. Commun. 11, 782 (2020).

32. Giuliano, C., Belin, D. & Everitt, B. J. Compulsive alcohol seeking results from a failure to disengage dorsolateral striatal control over behavior. J. Neurosci. 39, 1744–1754 (2019).

33. Brown, M. T. C. et al. Drug-driven AMPA receptor redistribution mimicked by selective dopamine neuron stimulation. PLoS One 5, e15870 (2010).

34. Bariselli, S., Miyazaki, N. L., Creed, M. C. & Kravitz, A. V. Orbitofrontal-striatal potentiation underlies cocaine-induced hyperactivity. Nat. Commun. 11, 3996 (2020).

35. Stachniak, T. J., Ghosh, A. & Sternson, S. M. Chemogenetic Synaptic Silencing of Neural Circuits Localizes a Hypothalamus→Midbrain Pathway for Feeding Behavior. Neuron 82, 797–808 (2014).

36. Nestler, E. J. & Lüscher, C. The Molecular Basis of Drug Addiction: Linking Epigenetic to Synaptic and Circuit Mechanisms. Neuron 102, 48–59 (2019). doi:10.1016/j.neuron.2019.01.016

37. Padoa-Schioppa, C. & Assad, J. A. Neurons in the orbitofrontal cortex encode economic value. Nature 441, 223–226 (2006).

38. Goldstein, R. Z. & Volkow, N. D. Dysfunction of the prefrontal cortex in addiction: Neuroimaging findings and clinical implications. Nat. Rev. Neurosci. 12, 652–669 (2011).

39. Hirokawa, J., Vaughan, A., Masset, P., Ott, T. & Kepecs, A. Frontal cortex neuron types categorically encode single decision variables. Nature 576, 446–451 (2019)

40. Masset, P., Ott, T., Lak, A., Hirokawa, J. & Kepecs, A. Behavior- and modality-general representation of confidence in orbitofrontal cortex. Cell 182, 112–116 (2020).

41. Schoenbaum, G., Chang, C. Y., Lucantonio, F. & Takahashi, Y. K. Thinking Outside the Box: Orbitofrontal Cortex, Imagination, and How We Can Treat Addiction. Neuropsychopharmacology 41, 2966–2976 (2016).

42. Takahashi, Y. K. et al. Neural Estimates of Imagined Outcomes in the Orbitofrontal Cortex Drive Behavior and Learning. Neuron 80, 507–518 (2013).

43. Belin, D., Berson, N., Balado, E., Piazza, P. V. & Deroche-Gamonet, V. High-novelty-preference rats are predisposed to compulsive cocaine self-administration. Neuropsychopharmacology 36, 569–579 (2011).

44. Wall, N. R. et al. Complementary Genetic Targeting and Monosynaptic Input Mapping Reveal Recruitment and Refinement of Distributed Corticostriatal Ensembles by Cocaine. Neuron 104, 916–930 (2019).

45. Fanous, S. et al. Role of orbitofrontal cortex neuronal ensembles in the expression of incubation of heroin craving. J. Neurosci. 32, 11600–11609 (2012).

46. Bonson KR. et al. Neural Systems and Cue-Induced Cocaine Craving. Neuropsychopharmacology 26, 376–386 (2002).

47. Volkow, N. D., Wang, G. J., Fowler, J. S., Tomasi, D. & Telang, F. Addiction: Beyond dopamine reward circuitry. Proc. Natl. Acad. Sci. U. S. A. 108, 15037–15042 (2011).

48. Lucantonio, F., Stalnaker, T. A., Shaham, Y., Niv, Y. & Schoenbaum, G. The impact of orbitofrontal dysfunction on cocaine addiction. Nat. Neurosci. 15, 358–366 (2012).

